# Offline memory replay in recurrent neuronal networks emerges from constraints on online dynamics

**DOI:** 10.1101/2021.10.27.466186

**Authors:** Aaron D. Milstein, Sarah Tran, Grace Ng, Ivan Soltesz

## Abstract

During spatial exploration, neural circuits in the hippocampus store memories of sequences of sensory events encountered in the environment. When sensory information is absent during “offline” resting periods, brief neuronal population bursts can “replay” sequences of activity that resemble bouts of sensory experience. These sequences can occur in either forward or reverse order, and can even include spatial trajectories that have not been experienced, but are consistent with the topology of the environment. The neural circuit mechanisms underlying this variable and flexible sequence generation are unknown. Here we demonstrate in a recurrent spiking network model of hippocampal area CA3 that experimental constraints on network dynamics such as population sparsity, stimulus selectivity, rhythmicity, and spike rate adaptation enable additional emergent properties, including variable offline memory replay. In an online stimulus-driven state, we observed the emergence of neuronal sequences that swept from representations of past to future stimuli on the timescale of the theta rhythm. In an offline state driven only by noise, the network generated both forward and reverse neuronal sequences, and recapitulated the experimental observation that offline memory replay events tend to include salient locations like the site of a reward. These results demonstrate that biological constraints on the dynamics of recurrent neural circuits are sufficient to enable memories of sensory events stored in the strengths of synaptic connections to be flexibly read out during rest and sleep, which is thought to be important for memory consolidation and planning of future behavior.

## Introduction

In mammals, the hippocampus is a brain region involved in the storage and recall of spatial and episodic memories. As an animal explores a spatial environment, different subpopulations of hippocampal neurons known as “place cells” are selectively activated at different positions in space, resulting in sequences of neuronal spiking that are on the seconds-long timescale of locomotor behavior (O’Keefe and Conway, 1978). The synchronous firing of these sparse neuronal ensembles is coordinated by population-wide oscillations referred to as theta (∼4-10 Hz) and gamma (∼30-100 Hz) rhythms (Colgin, 2016). Within each cycle of the theta rhythm (∼125 ms), the spiking of active neurons is organized into fast timescale sequences such that neurons selective for just-visited positions spike first, then neurons selective for the current position, and finally neurons selective for the next and future positions spike last in the sequence (Drieu and Zugaro, 2019; Foster and Wilson, 2007). These order-preserving fast timescale “theta sequences” are thought to be involved in planning and learning of event order through associative synaptic plasticity (Jensen et al., 1996; Kay et al., 2020).

When an animal stops running, theta and gamma oscillations decrease, and neuronal circuits in the hippocampus instead emit intermittent synchronous bursts of activity that are associated with high-frequency oscillatory activity detectable in local field potential recordings in hippocampal area CA1. These ∼100-200 ms long events are referred to as “sharp wave-ripples” (SWRs) (Colgin, 2016), and they occur during non-locomotor periods of quiet wakefulness, during reward consumption, and during slow-wave sleep, when sensory information about the spatial environment is reduced or absent. During SWRs, sparse subsets of neurons are co-activated, with a tendency for neurons that fire sequentially during exploratory behavior to also fire sequentially during SWRs, either in the same order, in reverse, or with a mixture of both directions (Davidson et al., 2009; Pfeiffer, 2020; Stella et al., 2019; Wu and Foster, 2014). The hippocampus can also activate sequences of place cells during SWRs that correspond to possible paths through the environment that have not actually been experienced, suggesting a possible role for offline sequence generation in behavioral planning (Gupta et al., 2010; Igata et al., 2021; Ólafsdóttir et al., 2015, 2018; Wu and Foster, 2014). Manipulations that disrupt neuronal activity during SWRs result in deficits in memory recall (Girardeau et al., 2009; Jadhav et al., 2012), supporting a hypothesis that offline reactivation of neuronal ensembles during SWRs is important for the maintenance and consolidation of long-term memories (Buzsáki, 1989; Joo and Frank, 2018).

Hippocampal SWRs are thought to be generated by the synchronous firing of subpopulations of neurons in the CA2 and CA3 regions of the hippocampus (Csicsvari et al., 2000; Oliva et al., 2016), which are characterized by substantial recurrent feedback connectivity (Duigou et al., 2014; Guzman et al., 2016; Okamoto and Ikegaya, 2019). Recurrent networks have long been appreciated for their ability to generate rich internal dynamics (Amit and Brunel, 1997), including oscillations (Ermentrout, 1992). It has also been shown that associative plasticity at recurrent connections between excitatory neurons can enable robust reconstruction of complete memory representations from incomplete or noisy sensory cues (Amit and Brunel, 1997; Griniasty et al., 1993; Guzman et al., 2016; Hopfield, 1982; Marr, 1971; Treves and Rolls, 1994). However, this “pattern completion” function of recurrent networks requires that similarly tuned neurons activate each other via strong synaptic connections, resulting in sustained self-activation, rather than sequential activation of neurons that are selective for distinct stimuli (Lisman et al., 2005; Pfeiffer and Foster, 2015; Renno-Costa et al., 2014). Previous work has shown that, in order for recurrent networks to generate sequential activity, some mechanism must be in place to “break the symmetry” and enable spread of activity from one ensemble of cells to another (Sompolinsky and Kanter, 1986; Tsodyks et al., 1996). During spatial navigation, feedforward sensory inputs carrying information about the changing environment can provide the momentum necessary for sequence generation. However, during hippocampal SWRs, sensory inputs are reduced, and activity patterns are thought to be primarily internally generated by the recurrent connections within the hippocampus. In this study we use a computational model of hippocampal area CA3 to investigate the synaptic, cellular, and network mechanisms that enable flexible offline generation of memory-related neuronal sequences in the absence of ordered sensory information.

A number of possible mechanisms for sequence generation in recurrent networks have been proposed:

1. Winner-take-all network mechanism (Almeida et al., 2007, 2009a; Lisman and Jensen, 2013): Within this framework, the subset of excitatory neurons receiving the most strongly weighted synaptic inputs responds first upon presentation of a stimulus. This active ensemble of cells then recruits feedback inhibition via local interneurons, which in turn prevents other neurons from firing for a brief time window (e.g. the ∼15 ms duration of a single gamma cycle). This highlights the important roles that inhibitory neurons play in regulating sparsity (how many cells are co-active), selectivity (which cells are active), and rhythmicity (when cells fire) in recurrent networks (Almeida et al., 2009b; Rennó-Costa et al., 2019; Stark et al., 2014; Stefanelli et al., 2016). However, while oscillatory feedback inhibition provides a network mechanism for parsing neuronal sequences into discrete elements, additional mechanisms are still required to ensure that distinct subsets of excitatory neurons are activated in a particular order across successive cycles of a rhythm (Lisman et al., 2005; Ramirez-Villegas et al., 2018).
2. Heterogeneous cellular excitability (Luczak et al., 2007; Stark et al., 2015): If the intrinsic properties of neurons in a network are variable and heterogeneous, when a stimulus is presented, those neurons that are the most excitable will fire early, while neurons with progressively lower excitability will fire later, resulting in sequence generation. This mechanism can explain the offline generation of stereotyped, unidirectional sequences, but cannot account for variable generation of sequences in both forward and reverse directions.
3. Asymmetric distributions of synaptic weights (Sompolinsky and Kanter, 1986; Tsodyks et al., 1996): During learning, if changes in synaptic weights are controlled by a temporally asymmetric learning rule, recurrent connections can become biased such that neurons activated early in a sequence have stronger connections onto neurons activated later in a sequence (Levy, 1989; Malerba and Bazhenov, 2019; McNaughton and Morris, 1987; Reifenstein et al., 2021). This enables internally generated activity to flow along the direction of the bias in synaptic weights. While this mechanism accounts for offline replay of specific sequences in the same order experienced during learning, it cannot account for reverse replay or the flexible generation of non-experienced sequences (Gupta et al., 2010; Igata et al., 2021; Ólafsdóttir et al., 2015, 2018; Wu and Foster, 2014).
4. Cellular or synaptic adaptation: It has also been proposed that short-term adaptation of either neuronal firing rate (Itskov et al., 2011; Treves, 2004) or synaptic efficacy (Romani and Tsodyks, 2015) can enable neuronal sequence generation in recurrent networks without asymmetric synaptic weights. According to this scheme, recently-activated neurons initially recruit connected partners with high efficacy, but continued spiking results in either a decrease in firing rate, or a decrease in the probability of neurotransmitter release. This causes connections to fatigue over time, and favors the sequential propagation of activity to more recently activated cells. These mechanisms do allow for the stochastic generation of neuronal sequences in either the forward or reverse direction, though they do not prescribe which or how many neurons will participate in a given replay event.

In this study, we sought to understand how neuronal sequence generation in hippocampal area CA3 depends on the structure and function of the underlying network. To do this, we constructed a computational neuronal network model comprised of recurrently connected excitatory and inhibitory spiking neurons, and tuned it to match experimental constraints on the spiking dynamics of CA3 during spatial navigation, including sparsity, selectivity, rhythmicity, and spike rate adaptation. We then analyzed the direction and content of neuronal sequences generated both “online” during simulated navigation, and “offline” during simulated rest. We found that when the network was driven by ordered sensory information in the online state, it generated forward-sweeping “theta sequences” that depended on the structure of recurrent connectivity in the network. In the offline state driven by noise, the network generated heterogenous memory replay events that moved either in forward, reverse, or mixed directions, and depended on network sparsity and rhythmicity, and neuronal stimulus selectivity and spike-rate adaptation. Model degeneracy analysis and network perturbations indicated that offline memory replay does not occur in networks with disrupted recurrent connectivity, or in networks lacking sparsity, selectivity, rhythmicity, or spike rate adaptation. Finally, when particular spatial locations were over-represented by the network, as occurs in the hippocampus at sites of reward (Lee et al., 2006; Turi et al., 2019; Zaremba et al., 2017), memory replay events were biased towards trajectories that included those salient positions (Gillespie et al., 2021; Ólafsdóttir et al., 2015; Singer and Frank, 2009).

## Results

To investigate how sequential activity in the hippocampus generated “online” during spatial exploration can be recapitulated “offline” in the absence of sensory cues, we constructed a simple spiking neuronal network model of rodent hippocampal area CA3 (Materials and Methods). Neural circuits in the hippocampus and cortex typically comprise a majority of excitatory neurons that project information to downstream circuits, and a minority of primarily locally connected inhibitory interneurons. We included populations of excitatory (1000) and inhibitory (200) neurons in proportion to experimental observations (Pelkey et al., 2017; Tremblay et al., 2016) (Figure 1A). Cell models were single-compartment, integrate-and-fire neurons with saturable, conductance-based excitatory and inhibitory synapses (Carnevale and Hines, 2006; Izhikevich, 2007; Izhikevich and Edelman, 2008). Excitatory neurons were endowed with spike-rate adaptation to support punctuated bursting behavior during theta oscillations (O’Keefe and Recce, 1993; Scharfman, 1993), and inhibitory neurons exhibited fast-spiking dynamics to sustain continuous high frequency firing during gamma oscillations (Csicsvari et al., 2003; Ylinen et al., 1995) (Supplementary Figure S1A, Materials and Methods).

**Figure 1.**
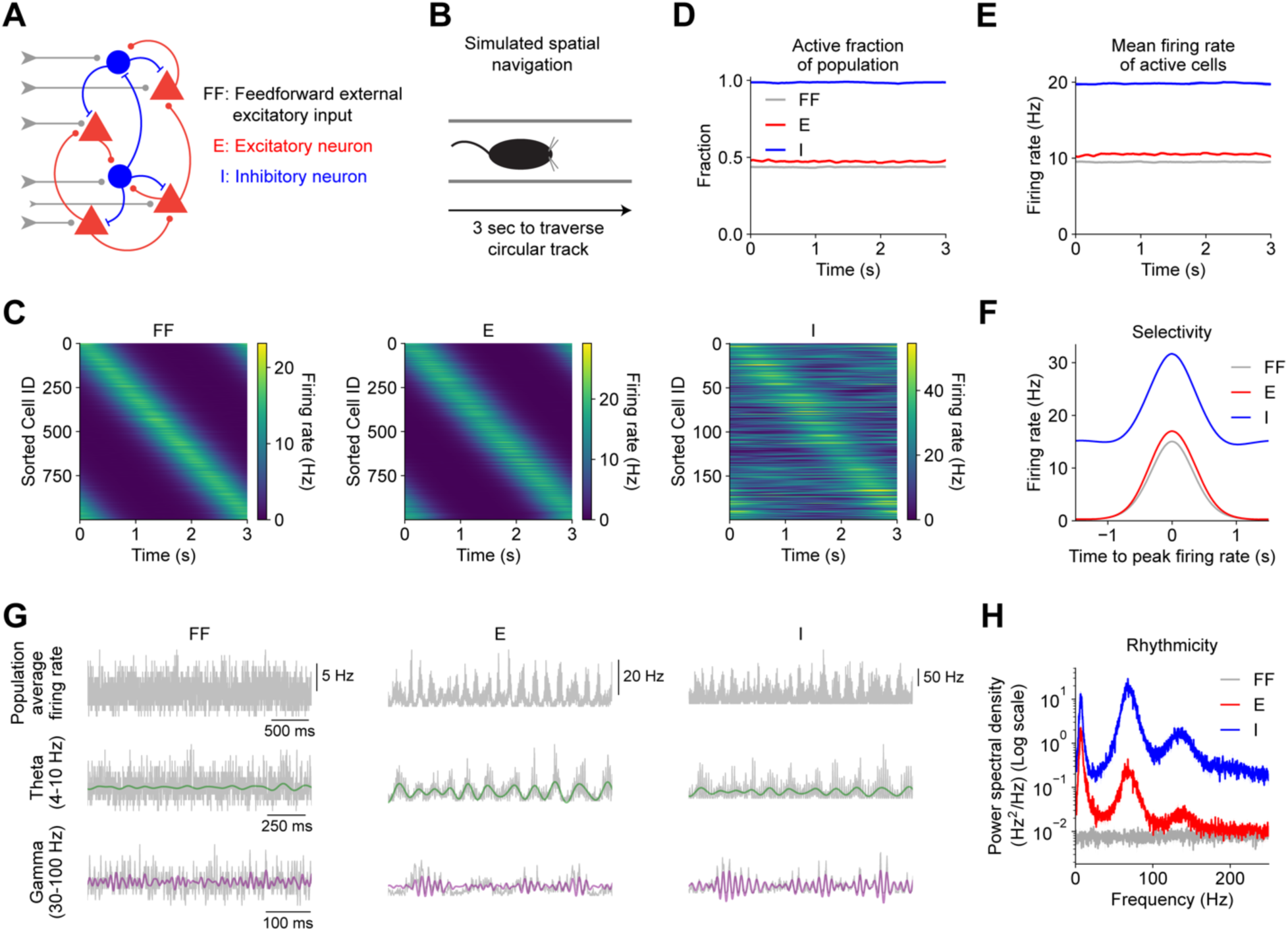
Sparsity, selectivity, and rhythmicity in a recurrent spiking neuronal network model of hippocampal area CA3. (**A**) Diagram illustrates connectivity of network model. Feedforward (FF) external excitatory inputs contact excitatory (E) and inhibitory (I) neurons. E and I neurons are recurrently connected to other E and I neurons. (**B**) Simulations of rodent “online exploration” emulated the response of the hippocampus during unidirectional locomotion along a circular linear track that takes 3 seconds to traverse at constant run velocity. (**C**) Population sparsity (active fraction of neurons) vs. time shown for each cell population. (**D**) Mean firing rate of active neurons vs. time shown for each cell population. (**E**) Firing rates vs. time of all neurons in each cell population are shown (average of 5 trials from one example network instance). Cells in each population are sorted by the location of maximum firing. (**F**) Average stimulus selectivity of each cell population. Trial-averaged activity of each cell was centered around the location of maximum firing, and then averaged across cells. (**G**) The average activity of each population on a single trial (top row) was bandpass filtered in the theta (middle row) and gamma (bottom row) frequency bands. Colored traces show filtered signals (theta: green, gamma: purple). Traces derived from one example network instance. (**H**) Power spectrum of average population activity indicates dominant frequency components in the theta and gamma bands (one-sided paired t-tests: theta: E vs. FF, p=0.00001; I vs. FF, p<0.00001; gamma: E vs. FF, p<0.00001; I vs. FF, p<0.00001). In (C), (D), (F), and (H), data were first averaged across 5 trials per network instance. Mean (solid) ± SEM (shading) were computed across 5 independent instances of each network model. p-values reflect FDR correction for multiple comparisons.

To simulate the sensory experience of locomotion in a spatial environment, we provided both excitatory and inhibitory neurons with external afferent inputs from a population of 1000 excitatory neurons, each of which was selectively activated at distinct but overlapping positions within a simulated circular track that took 3 seconds to traverse (Figures 1A – 1C, Materials and Methods). Recurrent connections within and between excitatory and inhibitory cell populations were also included (Figure 1A), as they are hallmark features of hippocampal area CA3, and have been shown to support rich network dynamics (Renno-Costa et al., 2014; Stark et al., 2014). Specifically, inhibitory feedback connections have been shown to regulate the number of simultaneously active neurons (sparsity) (Stefanelli et al., 2016), and to contribute to the generation of theta and gamma network oscillations (Bezaire et al., 2016; Geisler et al., 2005; Rennó-Costa et al., 2019; Stark et al., 2014; Wang, 2010). Plastic excitatory connections between excitatory neurons have long been implicated in stimulus selectivity and the storage and recall of memories (Almeida et al., 2007; Hopfield, 1982; Lisman and Jensen, 2013). It has been proposed that strong connections between ensembles of co-active neurons could arise through a combination of biased connectivity during brain development (Buzsáki et al., 2021; Dragoi and Tonegawa, 2013; Farooq and Dragoi, 2019; Grosmark and Buzsáki, 2016), and experience-driven synaptic plasticity during learning (Bittner et al., 2015, 2017; Brunel and Trullier, 1998; Káli and Dayan, 2000; Milstein et al., 2020; O’Neill et al., 2008). While here we did not simulate these dynamic processes explicitly, we implemented the structured connectivity that is the end result of these processes by increasing the strengths of synaptic connections between excitatory cells that share overlapping selectivity for spatial positions in the environment (Table 1, Supplementary Figure S1B, Materials and Methods) (Arkhipov et al., 2018).

**Table 1.** Model parameter values.

| Parameter | Bounds | Structured<br>E <- E<br>weights | Random<br>E <- E<br>weights | Shuffled<br>E <- E<br>weights | Suppressed<br>rhythmicity | No sparsity<br>or selectivity<br>constraints | No spike<br>rate<br>adaptation |
| --- | --- | --- | --- | --- | --- | --- | --- |
| E <- FF<br>weight<br>mean | 0.1 – 5 | 1.28 | 0.29 | 1.30 | 0.37 | 1.57 | 0.55 |
| E <- FF<br>weight st.<br>dev. | 0.1 – 5 | 1.09 | 0.21 | 0.91 | 0.80 | 0.88 | 0.41 |
| E <- E weight<br>mean | 0.1 – 5 | 0.75 | 0.50 | 1.43 | 0.90 | 0.54 | 2.45 |
| E <- E weight<br>st. dev. | 0.1 – 5 | 0.52 | 0.48 | 0.58 | 0.25 | 0.73 | 1.74 |
| E <- I weight<br>mean | 0.1 – 5 | 0.87 | 0.70 | 2.06 | 0.86 | 0.59 | 0.54 |
| E <- I weight<br>st. dev. | 0.1 – 5 | 0.47 | 0.32 | 0.41 | 0.76 | 0.73 | 0.16 |
| I <- FF weight<br>mean | 0.1 – 5 | 1.50 | 1.83 | 2.44 | 0.38 | 1.53 | 1.60 |
| I <- FF weight<br>st. dev. | 0.1 – 5 | 0.68 | 0.82 | 0.45 | 0.38 | 0.55 | 0.22 |
| I <- E weight<br>mean | 0.1 – 5 | 1.85 | 1.31 | 1.44 | 1.63 | 1.77 | 0.82 |
| I <- E weight<br>st. dev. | 0.1 – 5 | 0.22 | 0.28 | 0.21 | 0.17 | 0.14 | 0.79 |
| I <- I weight<br>mean | 0.1 – 5 | 0.16 | 1.07 | 0.26 | 2.51 | 0.15 | 0.23 |
| I <- I weight<br>st. dev. | 0.1 – 5 | 0.12 | 0.68 | 0.47 | 0.33 | 0.77 | 0.05 |
| E <- FF and<br>E <- E max<br>structured<br>Δweight | 1 - 5 | 3.63 | 2.80 | 4.42 | 3.91 | 3.63 | 4.00 |
| E <- FF and<br>E <- E decay<br>(ms) | 2 - 20 | 3.43 | 10.76 | 3.48 | 19.87 | 3.53 | 5.91 |
| E <- I decay<br>(ms) | 2 - 30 | 2.32 | 3.89 | 2.72 | 17.05 | 2.53 | 27.87 |
| I <- FF and<br>I <- E decay<br>(ms) | 2 - 20 | 15.04 | 18.53 | 17.94 | 13.49 | 17.10 | 18.39 |
| I <- I decay<br>(ms) | 2 - 30 | 8.09 | 5.65 | 8.12 | 9.24 | 10.19 | 22.35 |
| E <- FF #<br>synapses<br>/ pair | 0 - 2 | 0.62 | 0.96 | 0.70 | 0.57 | 0.73 | 0.18 |
| E <- E #<br>synapses<br>/ pair | 0 - 2 | 0.55 | 0.57 | 0.55 | 0.05 | 0.33 | 0.46 |
| E <- I #<br>synapses<br>/ pair | 0 - 10 | 5.12 | 3.36 | 4.75 | 7.93 | 5.71 | 7.79 |
| I <- FF #<br>synapses<br>/ pair | 0 - 2 | 0.26 | 0.77 | 0.19 | 0.21 | 0.19 | 0.12 |
| I <- E #<br>synapses<br>/ pair | 0 - 2 | 0.31 | 0.47 | 0.19 | 0.22 | 0.13 | 0.60 |
| I <- I #<br>synapses<br>/ pair | 0 - 10 | 8.17 | 4.81 | 9.07 | 8.83 | 9.43 | 7.38 |

Despite the relatively simple architecture of this network model, a wide range of networks with distinct dynamics could be produced by varying a number of parameters, including 1) the probabilities of connections between cell types (Káli and Dayan, 2000), 2) the kinetics and strengths of synaptic connections between cell types (Brunel and Wang, 2003), and 3) the magnitude of the above-mentioned increase in synaptic strengths between neurons with shared selectivity (Brunel, 2016; Dorkenwald et al., 2019). To calibrate the network model to produce dynamics that matched experimentally-derived targets, we performed an iterative stochastic search over these parameters, and optimized the following features of the activity of the model network: 1) population sparsity - the fraction of active neurons of each cell type, 2) the mean firing rates of active neurons of each cell type, 3) stimulus-selective firing of excitatory cells, and 4) the frequency and amplitude of theta and gamma oscillations in the synchronous spiking activity of each cell population (Materials and Methods).

This procedure identified a model with dynamics that met all of the above constraints. Given sparse and selective feedforward inputs during simulated navigation (Figures 1B and 1C), the excitatory neurons in the network responded with a fraction of active cells (Figure 1D) and with average firing rates comparable to the those of the feedforward input population (Figure 1E). The majority of inhibitory neurons were activated continuously (Figures 1C and 1D) at high firing rates (Figure 1E). While excitatory neurons received random connections from feedforward afferents and from other excitatory neurons with heterogeneous spatial tuning, excitatory cells exhibited a high degree of spatial selectivity (Figures 1C and 1F). This selective increase in firing rate at specific spatial locations within the “place field” of each excitatory neuron was supported by enhanced synaptic connection strengths between excitatory neurons with overlapping tuning (Supplementary Figure S1B). While substantial background excitation occurred in all cells at all spatial positions, firing outside the place field of each cell was suppressed by sufficiently strong inhibitory input (Bittner et al., 2015; Grienberger et al., 2017). Interestingly, inhibitory neurons also exhibited spatial selectivity, albeit to a weaker degree and with a higher background firing rate (Figures 1C and 1F). This feature of the network dynamics was an emergent property that was not explicitly designed or optimized. While excitatory connections onto inhibitory cells were random and not weighted according to shared selectivity (Supplementary Figure S1B), the total amount of excitatory input arriving onto individual inhibitory cells fluctuated across spatial positions, and predicted a small degree of spatial selectivity (Supplementary Figure S1C). Inhibitory inputs received by inhibitory cells reduced their average activity, effectively enabling fluctuations in excitation above the mean to stand out from the background excitation (Supplementary Figure S1C and S1D). This mechanism of background subtraction by inhibitory synaptic input may explain the partial spatial selectivity previously observed in subpopulations of hippocampal inhibitory neurons (Ego-Stengel and Wilson, 2007; Geiller et al., 2020; Grienberger et al., 2017; Hangya et al., 2010; Marshall et al., 2002; Wilent and Nitz, 2007).

The tuned network model also exhibited oscillatory synchrony in the firing of the excitatory and inhibitory neuron populations, despite being driven by an asynchronous external input (Figures 1G and 1H). The requirement that the network self-generate rhythmic activity in the theta band constrained recurrent excitatory connections to be relatively strong, as this input provided the only source of rhythmic excitation within the network (Supplementary Figure S1E). Interestingly, as the firing rates of inhibitory cells increased within each cycle of the theta rhythm, their synchrony in the gamma band increased, resulting in an amplitude modulation of gamma paced at the theta frequency (Figure 1G and Supplementary Figure S1F). This “theta-nested gamma” is a well-known feature of oscillations in the hippocampus (Soltesz and Deschenes, 1993; Ylinen et al., 1995), and here emerged from fundamental constraints on dual band rhythmicity without requiring additional mechanisms or tuning.

### Position decoding reveals “theta sequences” during simulated navigation

Next, we analyzed neuronal sequence generation within the network during simulated navigation. First, we simulated multiple trials and computed trial-averaged spatial firing rate maps for all neurons in the network (Figure 1C). We then used these rate maps to perform Bayesian decoding of spatial position given the spiking activity of all cells in the network from individual held-out trials not used in constructing the decoding template (Figure 2A, Materials and Methods) (Davidson et al., 2009; Zhang et al., 1998). For the population of feedforward excitatory inputs, the underlying spatial firing rates were imposed, and the spikes of each cell were generated by sampling from an inhomogeneous Poisson process. Thus, decoding position from the activity of this population served to validate our decoding method, and indeed simulated position could be decoded from the spiking activity of the feedforward input population with very low reconstruction error (Figures 2A and 2D). When we applied this method to the population of excitatory neurons within the network, reconstruction error was increased (Figures 2A and 2D). This reflected an increased fraction of temporal bins (20 ms) where the decoded position was either behind or in advance of the actual position (Figure 2A). However, rather than simply reflecting reconstruction noise or poor spatial selectivity of individual cells (Figure 1F), these divergences from actual position resulted from consistent sequential structure in the spiking activity of cells in the excitatory population (Figure 2A). Ordered neuronal firing resulted in decoded positions that continuously swept from past positions, through the current actual position, to future positions, and then reset to past positions, on the timescale of the ongoing theta rhythm. These “theta sequences” caused decoded position estimates to oscillate around the actual position (Figure 2A), and this theta timescale oscillation accounted for a large proportion of the variance in decoded position (Figure 2E, Materials and Methods). Interestingly, we found that position could also be accurately decoded from the moderately spatially-tuned activity of inhibitory cells in the network (Figures 2A and 2D), and that the spiking activity of the inhibitory population was also organized into theta sequences (Figures 2A and 2E).

**Figure 2.**
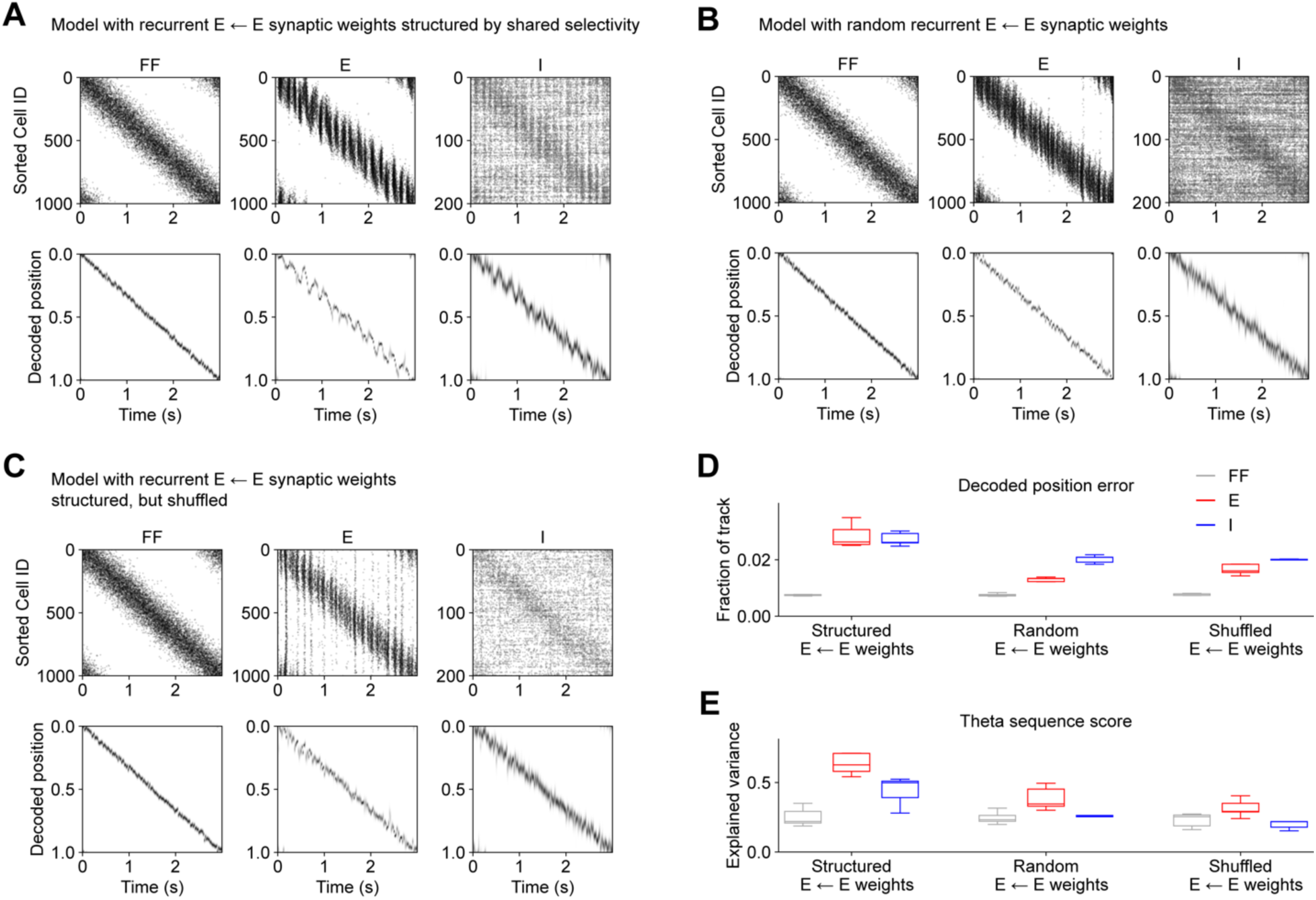
Online neuronal sequence generation depends on recurrent excitatory synaptic connectivity. (**A**) “Online exploration” was simulated for the same network model as in Figure 1, in which recurrent excitatory connections between E cells were structured such that neurons with shared selectivity have elevated synaptic weights. Top row: spike times of all neurons in each cell population on a single trial of simulated “online exploration” are marked. A separate set of 5 trials was used to construct a spatial firing rate template for each neuron (shown in Figure 1C). Cells in each population are sorted by the location of maximum average spatial firing rate. Bottom row: the spatial firing rate templates for all neurons were used to perform Bayesian decoding of spatial position from the single trial spiking data shown in the top row. For each cell population, the likelihood of each spatial position in each time bin (20 ms) is indicated by grayscale intensity. (**B**) Same as (A) for alternative network model with random synaptic strengths at recurrent excitatory connections between E cells. Spatial firing rate templates used for decoding are shown in Supplementary Figure S2B. (**C**) Same as (A) for alternative network model in which the structured excitatory recurrent synaptic weights between E cells were randomly shuffled. Spatial firing rate templates used for decoding are shown in Supplementary Figure S2H. (**D**) Decoded position error is quantified as the difference between actual and predicted position. The absolute value of decoded position error is expressed as a fraction of the track length (one-sided paired t-tests: Structured E ← E weights: E vs. FF, p<0.00001; I vs. FF, p<0.00001; Random E ← E weights: E vs. FF, p=0.00001; I vs. FF, p=0.00001; Shuffled E ← E weights: E vs. FF, p=0.00001; I vs. FF, p=0.00001; two-sided t-tests vs. data from model with structured E ← E weights: Random E ← E weights: E, p<0.00001, I, p=0.00001; Shuffled E ← E weights: E, p<0.00001, I, p=0.00001). (**E**) In the model with structured E ← E weights, decoded positions of E and I cell populations oscillated between past, current, and future positions at the timescale of the population theta oscillation. A theta sequence score was computed as the proportion of the variance in the decoded position error explained by a theta timescale oscillation (see Materials and Methods) (one-sided paired t-tests: Structured E ← E weights: E vs. FF, p=0.00005; I vs. FF, p-0.00005; Random E ← E weights: E vs. FF, p=0.00002; I vs. FF, p=0.28287; Shuffled E ← E weights: E vs. FF, p=0.00010; I vs. FF, p=0.99600; two-sided t-tests vs. data from model with structured E ← E weights: Random E ← E weights: E, p<0.00001, I, p<0.00001; Shuffled E ← E weights: E, p<0.00001, I, p<0.00001). In (D) and (E), data were first averaged across 5 trials per network instance. Box and whisker plots depict data from 5 independent instances of each network model (see Materials and Methods). p-values reflect FDR correction for multiple comparisons.

A number of possible mechanisms have been proposed to account for theta sequence generation *in vivo*, including synaptic, cell-intrinsic, and network-level mechanisms (Chadwick et al., 2015, 2016; Drieu and Zugaro, 2019; Foster and Wilson, 2007; Grienberger et al., 2017; Kang and DeWeese, 2019; Mehta et al., 2002; Skaggs et al., 1996). That theta sequences in the model emerged in both excitatory and inhibitory neuron populations implicates recurrent interactions within the network (Chadwick et al., 2016). To further investigate, we analyzed neuronal sequence generation in a variant of the model in which the strengths of recurrent connections between excitatory neurons were randomized and no longer depended on shared spatial selectivity between connected pairs of cells (Supplementary Figure S2A). This alternative model could still be tuned to match experimental targets, including sparsity, selectivity, and rhythmicity (Supplementary Figures S2B – S2F). In this case the spatial selectivity of excitatory cells was entirely determined by the synaptic weights of the feedforward afferent inputs (Supplementary Figure S2A), while the recurrent excitatory input supported synchronization in the theta and gamma bands (Supplementary Figure S2F). However, in this model, theta timescale neuronal sequence generation in both excitatory and inhibitory cells was suppressed (Figures 2B and 2E). Decoding of position from spikes on single trials produced lower reconstruction error (Figure 2D), as neuronal population activity more faithfully followed the current spatial position provided by the feedforward inputs, and was not organized into the sweeps from past to future positions characteristic of theta sequences (Figures 2B and 2E). We also tested a related variant of the model in which the skewed distribution of recurrent excitatory synaptic weights used in the structured weights model (Figures 1 and 2A, and Supplementary Figure 1) was randomly shuffled (Figure 2C and Supplementary Figures S2G – S2L). Theta sequence generation was also reduced in this network model variant (Figure 2E). These results indicate that structure in the synaptic strengths of recurrent excitatory connections is required for the generation of fast timescale (∼125 ms) neuronal sequences when network activity is driven by behavioral timescale (> 1 s) sequences of sensory inputs, as occurs during spatial exploration.

### Emergence of offline memory replay

The above results show that the same network structure that enables population dynamics in CA3 to exhibit sparsity, selectivity, and rhythmicity also supports neuronal sequence generation in the online state when ordered sensory information is present. We next sought to understand how neuronal sequences consistent with the sensory environment are generated offline when sensory inputs are reduced. To mimic the transient (∼50-150 ms) increase in population activity that occurs during a hippocampal SWR (Fernández-Ruiz et al., 2019), we transiently stimulated the network by randomly choosing sparse subsets of cells from the feedforward input population to emit spike (Figures 3A and 3B). We then used the same decoding templates as above, constructed from the trial-averaged activity during simulated run, to decode spatial position from spiking activity during these transient offline events (Materials and Methods).

**Figure 3.**
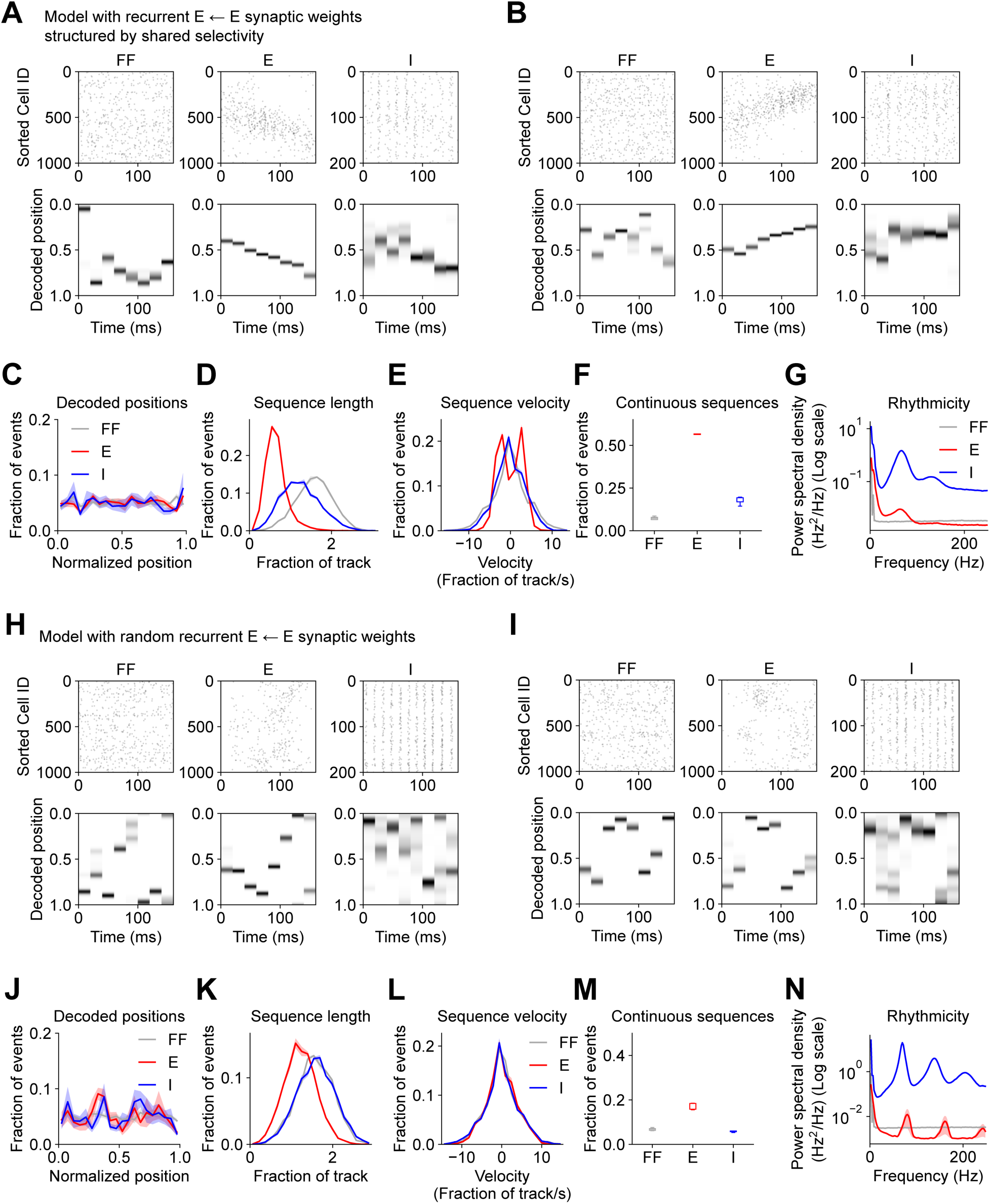
Forward and reverse offline memory replay depends on recurrent excitatory synaptic connectivity. (**A** – **B**) “Offline rest” was simulated for the network model with structured E ← E weights (Figure 1). Top row: spike times of all neurons in each cell population on a single trial of simulated “offline rest” are marked. Data from 5 trials of simulated “online exploration” was used to construct a spatial firing rate template for each neuron (Figure 1C). Cells in each population are sorted by the location of maximum average spatial firing rate. Bottom row: the spatial firing rate templates for all neurons were used to perform Bayesian decoding of spatial position from the single trial spiking data shown in the top row. For each cell population, the likelihood of each spatial position in each time bin (20 ms) is indicated by grayscale intensity. (A) and (B) correspond to two example trials from one example network instance. (**C** – **G**) This procedure was repeated for 1000 trials for each of 5 instances of the network. (**C**) Histogram of spatial positions decoded from each cell population across all simulated replay events (two-sided K-S tests: E vs. FF, p=0.99974; I vs. FF, p=0.99974). (**D**) Histogram of the path length of spatial sequences decoded from each cell population (two-sided K-S tests: E vs. FF, p<0.00001; I vs. FF, p<0.00001). (**E**) Histogram of the mean velocity of spatial sequences decoded from each cell population (two-sided K-S tests: E vs. FF, p=0.00235; I vs. FF, p=0.73515). (**F**) Fraction of events that met criterion for sequences consistent with continuous spatial trajectories (see Materials and Methods) (one-sided paired t-tests: E vs. FF, p<0.00001, I vs. FF, p=0.00084). (**G**) Power spectrum of average population activity indicates high frequency components (one-sided paired t-tests: 75 Hz – 300 Hz frequency band: E vs. FF, p=1; I vs. FF, p<0.00001). (**H** – **N**) Same as (A – G) for an alternative network model with random E ← E weights. (**H**) and (**I**) correspond to two example trials from one example network instance. (**J**) Decoded positions (two-sided K-S tests: E vs. FF, p=0.99974; I vs. FF, p=0.99974; two-sided K-S tests vs. data from model with structured E ← E weights in (C): E, p=0.85869; I, p=0.87577). (**K**) Offline sequence path length (two-sided K-S tests: E vs. FF, p<0.00001; I vs. FF, p=0.99997; two-sided K-S tests vs. data from model with structured E ← E weights in (D): E, p<0.00001; I, p<0.00001). (**L**) Offline sequence velocity (two-sided K-S tests: E vs. FF, p=0.99877; I vs. FF, p=0.99877; two-sided K-S tests vs. data from model with structured E ← E weights in (E): E, p=0.01591; I, p=0.76837). (**M**) Fraction of events that met criterion for sequences consistent with continuous spatial trajectories (see Materials and Methods) (one-sided paired t-tests: E vs. FF, p=0.00005, I vs. FF, p=0.86564; two-sided t-tests vs. data from model with structured E ← E weights in (F): E, p<0.00001; I, p=0.00001). (**N**) Offline high-frequency rhythmicity (one-sided paired t-tests: 75 Hz – 300 Hz frequency band: E vs. FF, p=0.97708; I vs. FF, p<0.00001; two-sided t-tests vs. data from model with structured E ← E weights in (G): E, p=0.64948; I, p<0.00001). In (C – E), (G), (J – L), and (N), mean (solid) ± SEM (shading) were computed across 5 independent instances of each network. In (F) and (M), box and whisker plots depict data from 5 independent instances of each network model (see Materials and Methods). p-values reflect FDR correction for multiple comparisons.

Given that the place field locations of the stimulated neurons in the feedforward input population were heterogeneous and unordered, the spatial positions decoded from their spiking were typically discontiguous across adjacent temporal bins (Figures 3A, 3B and 3F). This input pattern evoked spiking in sparse subsets of both the excitatory and inhibitory populations in the network (Figures 3A and 3B). In contrast with the feedforward population, the activity evoked in excitatory neurons was structured such that neurons with nearby place field locations spiked in adjacent temporal bins, resulting in decoded spatial trajectories that were continuous (Figures 3A, 3B and 3F). Inhibitory neuron activity during these events was organized into high-frequency oscillations (Figures 3A, 3B and 3G). This procedure was repeated to produce thousands of offline events evoked by stimulation of different random ensembles of inputs (Materials and Methods). Across these events, each position along the track was decoded with equal probability (Figure 3C). For each event, the length and mean velocity of the decoded trajectory was calculated from the differences in decoded positions between adjacent bins (Figures 3D and 3E). A mean velocity of zero corresponds to events with equal steps in the forward and reverse directions, while positive velocities correspond to net forward-moving trajectories, and negative velocities correspond to net backwards-moving trajectories. While the trajectories decoded from the random feedforward input population were comprised of large, discontiguous steps that traced out large path lengths with an average velocity near zero, the excitatory neuron population generated shorter, more continuous sequences that progressed in either the forward or reverse directions (Figures 3D – 3F). These trajectories on average covered ∼0.5 the length of the track in the short (∼150 ms) duration of the offline event. Compared to the run trajectory, which took 3 seconds to cover the full track length, this corresponded to a ∼10-fold temporal compression (Figure 3E), similar to experimental data (Davidson et al., 2009). Spatial trajectories decoded from the inhibitory neuron population were intermediate in length, but with little forward or reverse momentum, similar to the feedforward inputs. However, the inhibitory cells exhibited high-frequency synchrony (Figures 3A, 3B and 3G), similar to experimentally recorded CA3 interneurons during hippocampal SWRs (Csicsvari et al., 2000; Tukker et al., 2013).

These data demonstrate that random, unstructured input can evoke sequential activity in a CA3-like recurrent spiking network, with sequences corresponding to forward, reverse, or mixed direction trajectories through an experienced spatial environment. This self-generated memory-related activity implicates information stored in the synaptic weights of the recurrent connections within the network as being important for offline replay of experience. However, in most previous models, sequence generation was unidirectional, and was enabled by an asymmetric bias in the strengths of recurrent connections such that neurons encoding positions early in a sequence formed stronger synapses onto neurons encoding later positions (Levy, 1989; Malerba and Bazhenov, 2019; McNaughton and Morris, 1987; Reifenstein et al., 2021; Sompolinsky and Kanter, 1986; Tsodyks et al., 1996). In contrast, the current network flexibly generated sequences in forward, reverse, or mixed directions, and had symmetric recurrent connections such that synaptic strengths between pairs of excitatory neurons depend only on overlapping selectivity, not on sequence order. Does sequence generation in the present network still depend on recurrent connectivity? To test this, we first verified that including an asymmetric bias in the strengths of excitatory connections produced offline replay events that were biased towards forward sequences (Supplementary Figure S3). We next analyzed the sequence content of offline events generated in the variants of the network model with random (Figures 3H – 3N and Supplementary Figures S2A – S2F) or shuffled Supplementary Figures S2G – S2L and S4) recurrent connection weights. Indeed, without structure in the recurrent connection weights, spatial trajectories decoded from the activity of excitatory neurons was more similar to those of the feedforward inputs, consisting of large, discontinuous steps without forward or reverse momentum (Figures 3H – 3M and Supplementary Figures S4A – S4F). Still, these networks exhibited high-frequency oscillatory synchrony during these offline events (Figure 3N and Supplementary Figure S4G).

### Exploration of model diversity and degeneracy

The above results strongly supported the hypothesis that recurrent connectivity is important for offline sequence generation. During optimization of each of the alternative network model configurations shown above to meet the multiple objectives of sparsity, selectivity, and rhythmicity, we evaluated 30,000 variants of each model with different parameters (Materials and Methods). For each model configuration, the model with the lowest overall multi-objective error was chosen as the “best” model for further analysis, as shown in Figures 2 and 3 and Supplementary Figures S1, S2 and S4. However, we noted that the parameter values that specified these “best” models were variable across the multiple network configurations (Table 1). This raised the possibility that while many models with diverse parameters may produce networks with similar online dynamics (referred to as model degeneracy (Marder and Taylor, 2011)), perhaps only a smaller subset of models would additionally support the emergence of offline sequence generation. The fact that sequence generation was not observed for either of the “best” models with disrupted recurrent excitatory synaptic weights could reflect an incomplete sampling of the parameter space and a failure to identify models that both meet the objective criteria for online dynamics and produce offline sequences.

Therefore, in order to explore the diversity and degeneracy of models evaluated during optimization, we devised a method to identify models that performed similarly with respect to multiple optimization objectives, but were specified by divergent sets of parameters (Materials and Methods). For each model configuration, all model variants evaluated during parameter optimization were sorted by their Euclidean distance from the “best” model in the space of model parameters. This resulted in an error landscape (e.g. Figure 4A) in which models with similar parameters resulted in similar multi-objective error scores. We then identified models located at local minima in this error landscape, which formed a group of models that were distant from each other in parameter space, but similar to each other in terms of overall multi-objective error. We termed a set of such models as a “Marder group” after pioneering work characterizing degeneracy in biological systems (Marder and Taylor, 2011). For each alternative network model configuration, we selected the 5 members of this “Marder group” with the lowest multi-objective error (labeled “best” and “M1” – “M4” in Figure 4A), and evaluated their network dynamics during simulations of both online exploration and offline rest. We first verified that for all model configurations with and without structured recurrent excitatory connectivity analyzed above (Figures 1 – 3 and Supplementary Figures S1, S2 and S4), model variants within a “Marder group” exhibited considerable diversity across model parameters (Figures 4B and 4C), and met all objective criteria for population sparsity (Figure 4D), neuronal stimulus selectivity (Figure 4E), and rhythmogenesis in the theta and gamma bands (Figures 4F and 4G). However, only model variants with synaptic weights structured by shared stimulus selectivity exhibited theta sequences during online run (Figure 4H) and generated offline sequences consistent with continuous spatial trajectories (Figure 4I). This analysis demonstrated that generation of memory-related neuronal sequences by recurrent networks requires that information about the topology of the sensory environment is stored in the strengths of recurrent excitatory connections between excitatory neurons.

**Figure 4.**
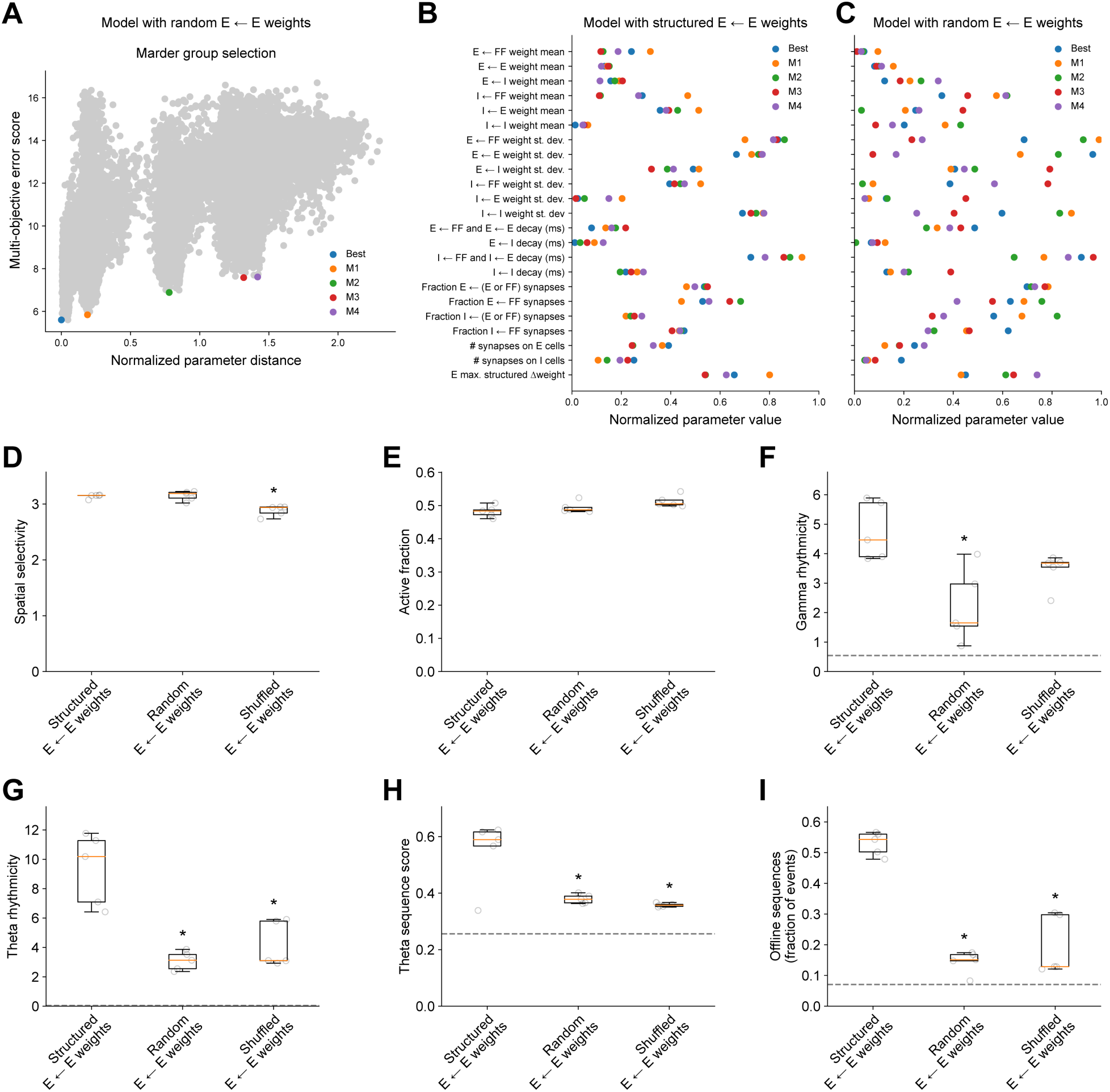
Exploration of model parameter diversity and degeneracy. (**A**) During network model optimization, 30,000 model variants with different parameters were evaluated. To explore model diversity and degeneracy, for each network model configuration, a subset of model variants termed a “Marder group” were selected based on their large distance from each other in the space of parameters, but their similar performance with respect to multiple optimization objectives (see Materials and Methods). This selection procedure is illustrated here for the model with random E ← E weights as an example. The 5 “Marder group” members with the lowest multi-objective error (labeled “best” and “M1” – “M4”) were selected for further evaluation. (**B**) For the network model configuration with structured E ← E weights, the range of parameter values across 5 distinct “Marder group” models are shown. (**C**) Same as (B) for the model with random E ← E weights. In (**D** – **I**), features of the simulated network dynamics produced by distinct model variants within a “Marder group” are compared across network model configurations. Each data point (grey circles) depicts one “Marder group” model. (**D**) Spatial selectivity of the excitatory neuron population during simulated “online exploration” is computed as a ratio of maximum to mean activity (two-sided t-tests vs. data from model with structured E ← E weights: Random E ← E weights: p=1; Shuffled E ← E weights: p=0.00224). (**E**) The fraction of the excitatory neuron population that is synchronously active during simulated “online exploration” is shown (two-sided t-tests vs. data from model with structured E ← E weights: Random E ← E weights: p=1; Shuffled E ← E weights: p=0.12079). (**F**) Gamma rhythmicity of the excitatory neuron population is computed as the integrated power spectral density in the gamma frequency band (two-sided t-tests vs. data from model with structured E ← E weights: Random E ← E weights: p=0.03552; Shuffled E ← E weights: p=0.16364). (**G**) Theta rhythmicity of the excitatory neuron population is computed as the integrated power spectral density in the theta frequency band (two-sided t-tests vs. data from model with structured E ← E weights: Random E ← E weights: p=0.00272; Shuffled E ← E weights: p=0.01944). (**H**) Theta sequence score (see Figure 2 and Materials and Methods) (two-sided t-tests vs. data from model with structured E← E weights: Random E ← E weights: p=0.02817; Shuffled E ← E weights: p=0.01473). (**I**) Fraction of events during simulated “offline rest” that met criterion for sequences consistent with continuous spatial trajectories (see Figure 3, Supplementary Figure S3, and Materials and Methods) (two-sided t-tests vs. data from model with structured E ← E weights: Random E ← E weights: p<0.00001; Shuffled E ← E weights: p=0.00044). In (D – I), for each network model configuration, box and whisker plots depict 5 distinct “Marder group” model variants with different parameters (see Materials and Methods). Data for each model variant (grey circles) were first averaged across 5 independent instances of that model variant. In (F – I), grey dashed lines indicate value for the feedforward input to the network for reference. p-values reflect Bonferroni correction for multiple comparisons.

### Constraints on online sparsity, selectivity and rhythmicity enable offline memory replay

Our above findings suggest that experimental constraints on the online dynamics of hippocampal area CA3 during spatial exploration are sufficient to enable the emergence of offline memory replay. We next sought to determine whether all or only a subset of these constraints were required for generation of memory-related sequences. To determine the importance of rhythmicity, we removed the optimization criteria that excitatory and inhibitory neuron populations synchronize in the theta and gamma bands, and instead added an objective to minimize power density across the full frequency spectrum (Supplementary Figure S5E). Following optimization, this alternative network model exhibited reduced rhythmicity, but still met objectives related to sparsity and selectivity (Supplementary Figures S5A – S5D). However, when challenged with random stimuli during simulated offline rest, this model with suppressed rhythmicity failed to generate continuous forward or reverse sequences (Supplementary Figures S5F – S5L). Rather, spatial trajectories decoded from offline population activity contained large discontiguous jumps in position, and most events had zero net velocity in either the forward or reverse direction. This indicated that the same reciprocal interactions between excitatory and inhibitory neurons that support rhythmogenesis in the theta and gamma bands also contribute to the sequential organization of neuronal activity during offline memory replay.

We next optimized a network model variant without constraints on population sparsity and neuronal stimulus selectivity (see Materials and Methods). In this network model, while feedforward excitatory inputs remained spatially tuned, their connectivity with excitatory neurons was shuffled to prevent inheritance of spatial selectivity. This resulted in a complete loss of sparsity of excitatory neuron activity (Supplementary Figures S6A and S6B), and suppressed stimulus selectivity in excitatory neurons even below the level exhibited in the inhibitory neuron population (Supplementary Figures S6C and S6D). Rhythmogenesis in the theta and gamma bands in excitatory and inhibitory neurons was maintained (Supplementary Figure S6E). During simulation of offline rest, this network generated highly synchronous population bursts that tended to either hover at one decoded position, or make large discontiguous jumps between positions (Supplementary Figure S6F – S6K). These results suggest that the network connectivity parameters that support highly sparse and selective neuronal activity in the online stimulus-driven state, also enable sparse reactivation of neuronal sequences in the offline state. We also repeated the model degeneracy analysis described above (Figure 4) for multiple model variants with compromised sparsity, selectivity or rhythmicity, which corroborated these findings (Supplementary Figure S8).

### Role of neuronal spike rate adaptation in forward and reverse offline memory replay

Above we showed that structure in the synaptic connectivity of the CA3 network is important for neuronal sequence generation. However, unlike previous models of sequence generation where asymmetry in connection strengths biased the direction of neuronal sequences (Levy, 1989; Malerba and Bazhenov, 2019; McNaughton and Morris, 1987; Reifenstein et al., 2021; Sompolinsky and Kanter, 1986; Tsodyks et al., 1996), here synaptic connectivity was symmetric, and yet variable neuronal sequences were flexibly generated in both forward and reverse directions (Figure 3A, 3B and 3E). If this symmetric connectivity enables recurrent networks to generate either forward or backward steps, what “breaks” this symmetry and produces sequences that make net progress in either the forward or backward direction? We next wondered if directionality of offline sequences in our network model was facilitated by our choice of “bursty” excitatory cell model, which exhibited spike-rate adaptation (Figure 5A). As mentioned before, use-dependent decreases in either firing rate or synaptic transmission over time can provide momentum to neuronal sequences by favoring the recruitment of new neurons that have not yet been activated during a network population event (Itskov et al., 2011; Romani and Tsodyks, 2015; Treves, 2004). To test a possible role for cellular adaptation in sequence generation in our model network, we replaced the “bursting” excitatory cell model with a “regular spiking” model without spike rate adaptation (Figure 5A). This cell model did not support the high instantaneous firing rates of the bursting cell model, which compromised the peak firing rates of excitatory cells in the network and their entrainment by higher frequency gamma oscillations during simulated online navigation (Supplementary Figure S7).

**Figure 5.**
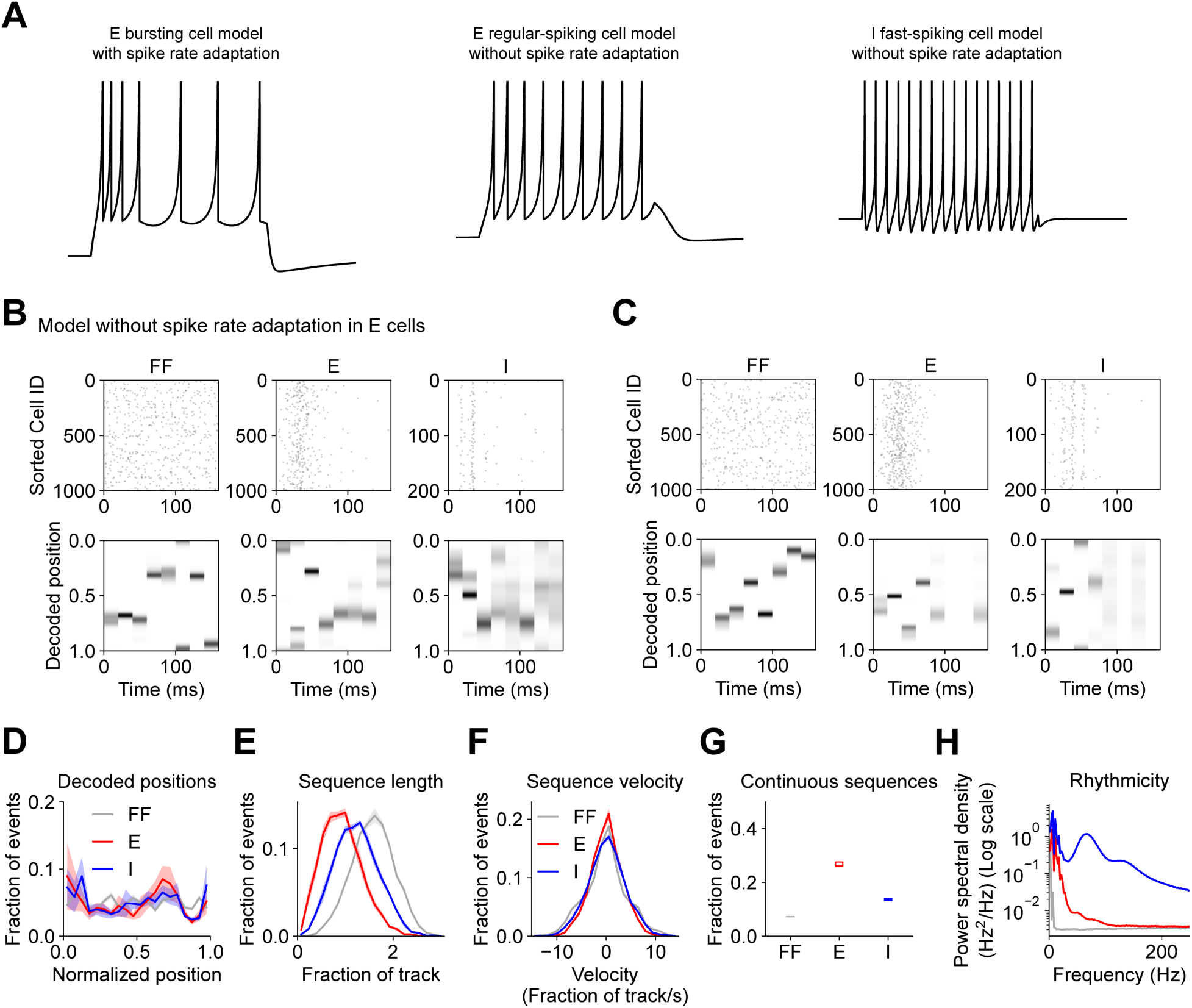
Neuronal spike rate adaptation supports offline memory replay. (**A**) Intracellular voltage recordings of three neuronal cell models with distinct spiking dynamics in response to simulated square-shaped intracellular current injections. (**B** – **H**) Same as Figures 3A – 2G for an alternative network model in which E cells are regular-spiking cell models without spike rate adaptation. (**B**) and (**C**) correspond to two example trials from one example network instance. (**D**) Decoded positions (two-sided K-S tests: E vs. FF, p=0.99974; I vs. FF, p=0.89825; two-sided K-S tests vs. data from model with structured E ← E weights in Figure 3C: E, p=0.84875; I, p=0.85869). (**E**) Offline sequence length (two-sided K-S tests: E vs. FF, p<0.00001; I vs. FF, p<0.00001; two-sided K-S tests vs. data from model with structured E ← E weights in Figure 3D: E, p<0.00001; I, p=0.99997). (**F**) Offline sequence velocity (two-sided K-S tests: E vs. FF, p=0.15282; I vs. FF, p=0.73515; two-sided K-S tests vs. data from model with structured E ← E weights in Figure 3E: E, p=0.03099; I, p=0.91017). (**G**) Fraction of events that met criterion for sequences consistent with continuous spatial trajectories (see Materials and Methods) (one-sided paired t-tests: E vs. FF, p=0.00038, I vs. FF, p=0.00119; two-sided t-tests vs. data from model with structured E ← E weights in Figure 3F: E, p<0.00001; I, p=0.03860). (**H**) Offline high-frequency rhythmicity (one-sided paired t-tests: 75 Hz – 300 Hz frequency band: E vs. FF, p=0.00114; I vs. FF, p=0.00003; two-sided t-tests vs. data from model with structured E ← E weights in Figure 3G: E, p<0.00001; I, p=0.00027). In (D – F) and (H), mean (solid) ± SEM (shading) were computed across 5 independent instances of each network. In (G), box and whisker plots depict data from 5 independent instances of each network model (see Materials and Methods). p-values reflect FDR correction for multiple comparisons.

Otherwise, this variant of the network did meet criterion for sparsity, selectivity, and rhythmicity (Supplementary Figures S7 and S8A – S8D). However, during simulated offline rest, random feedforward inputs evoked a truncated response from the network (Figures 5B and 5C), with the high frequency rhythmic activity of the inhibitory neurons diminishing before the end of the stimulus period (Figures 5B, 5C and 5H). Spatial trajectories decoded from the activity of excitatory neurons in the network were comprised of large steps that did not progress in either forward or reverse directions, similar to the random feedforward inputs (Figures 5E – 5G and Supplementary Figure S8E). These data show that adaptation of neuronal spiking provides a cellular-level mechanism for flexible and reversible sequence generation in recurrent spiking networks.

### Preferential replay of reward location

Thus far, we have simulated network activity during spatial navigation, and identified features of the network that enable offline replay of behavioral sequences stored in memory. However, in these simulations all spatial positions were visited with equal occupancy, and considered to be of equal salience or relevance to the virtual animal. This resulted in all positions being replayed with equal probability offline (Figure 3C), mimicking experimental conditions where all spatial positions contain discriminable sensory cues, and opportunities for reward are provided at random times and positions (Turi et al., 2019; Zaremba et al., 2017). However, it has been shown that when reward is provided at a fixed location that the animal must discover through learning, offline memory replay events become biased towards sequences of place cells that encode positions nearby and including the site of reward (Gillespie et al., 2021; Ólafsdóttir et al., 2018; Pfeiffer, 2020; Singer and Frank, 2009). In parallel with the development of this bias in offline memory replay during learning, it has been shown that the fraction of hippocampal pyramidal cells that selectively fire along the path to reward increases (Lee et al., 2006; Turi et al., 2019; Zaremba et al., 2017). Here we sought to test the hypothesis that this network-level over-representation of reward location is sufficient to bias the content of offline memory replay.

We chose a position along the virtual track to be the fixed location of a simulated reward, and biased the allocation of place field locations such that an increased fraction of excitatory neurons were selectively activated at positions near the reward (Figure 6A). As before, feedforward and recurrent synaptic connection strengths were increased between neurons with overlapping selectivity (Supplementary Figure S9A). Aside from an enhanced fraction of active excitatory neurons near the reward site (Supplementary Figure S9B), this produced network dynamics during simulated navigation that conformed to experimental constraints for sparsity, selectivity, and rhythmicity (Figure 6A and Supplementary Figures S9B – S9E). During simulated offline rest, the excitatory neurons in the network responded to random feedforward inputs by generating neuronal sequences corresponding to forward, reverse, and mixed direction trajectories through the environment (Figures 6B – 6G), paced by high frequency oscillations in the inhibitory cells (Figure 6H), as before (Figures 3A – 3G). However, now positions near the simulated reward site were replayed in a higher proportion of replay events (Figure 6D). This preferential replay of locations over-represented by the network recapitulated experimental findings and supported the hypothesis that nonuniform place cell allocation and biased memory replay are causally linked (Levy, 1989).

**Figure 6.**
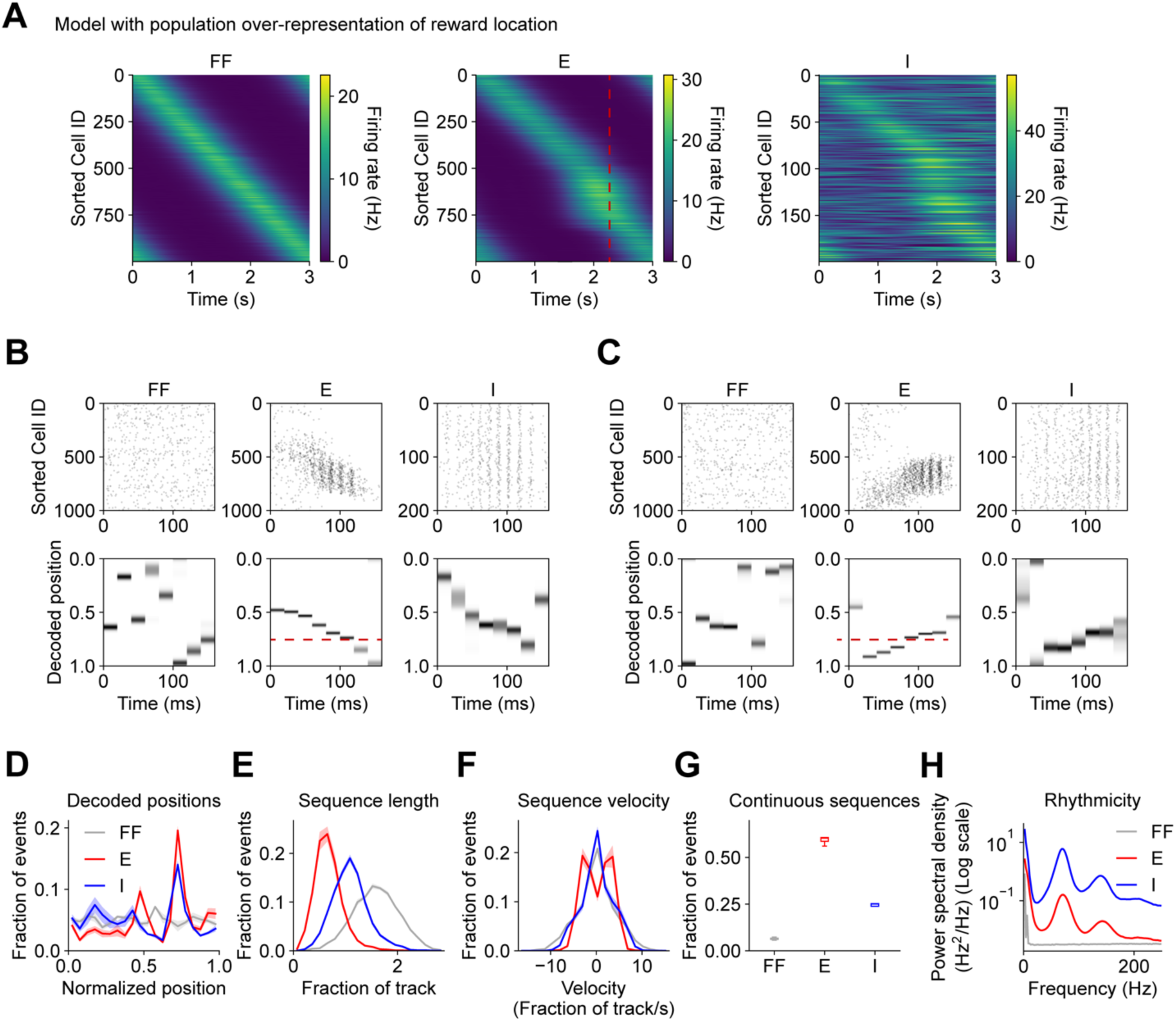
Offline memory replay is biased towards reward positions over-represented by the network. (**A**) In this variant of the network, an increased proportion of E cells are selective for spatial positions near the site of a simulated reward. Firing rates vs. time of all neurons in each cell population are shown (average of 5 trials from one example network instance). Cells in each population are sorted by the location of maximum firing. The simulated reward site is marked with red dashed line. (**B** – **H**) Same as Figures 3A – 2G for an alternative network model with population-level over-representation of reward location in E cells. (**B**) and (**C**) correspond to two example trials from one example network instance. The simulated reward site is marked with red dashed line. (**D**) Decoded positions (two-sided K-S tests: E vs. FF, p<0.00001; I vs. FF, p=0.41580; two-sided K-S tests vs. data from model with structured E ← E weights in Figure 3C: E, p<0.00001; I, p=0.84875). (**E**) Offline sequence length (two-sided K-S tests: E vs. FF, p<0.00001; I vs. FF, p<0.00001; two-sided K-S tests vs. data from model with structured E ← E weights in Figure 3D: E, p=0.25150; I, p<0.00001). (**F**) Offline sequence velocity (two-sided K-S tests: E vs. FF, p=0.02742; I vs. FF, p=0.73515; two-sided K-S tests vs. data from model with structured E ← E weights in Figure 3E: E, p=0.16847; I, p=0.62023). (**G**) Fraction of events that met criterion for sequences consistent with continuous spatial trajectories (see Materials and Methods) (one-sided paired t-tests: E vs. FF, p<0.00001, I vs. FF, p=0.00027; two-sided t-tests vs. data from model with structured E ← E weights in Figure 3F: E, p=0.05397; I, p=0.00754). (**H**) Offline high-frequency rhythmicity (one-sided paired t-tests: 75 Hz – 300 Hz frequency band: E vs. FF, p=0.00067; I vs. FF, p=0.00006; two-sided t-tests vs. data from model with structured E ← E weights in Figure 3G: E, p=0.00003; I, p=0.00001). In (D – F) and (H), mean (solid) ± SEM (shading) were computed across 5 independent instances of each network. In (G), box and whisker plots depict data from 5 independent instances of each network model (see Materials and Methods). p-values reflect FDR correction for multiple comparisons.

## Discussion

In this study we used a simple recurrent spiking network model of hippocampal area CA3 to investigate the structural and functional requirements for offline replay of spatial memories. We optimized synaptic, cellular, and network parameters of the network to produce population dynamics that match experimentally observed sparsity, selectivity and rhythmicity. We found that networks that fit these constraints exhibit additional emergent properties, including the ability to generate fast timescale memory-related neuronal sequences. During simulated spatial navigation, when ordered sensory information was provided on the seconds-long timescale of locomotion behavior, the network produced neuronal sequences that swept from past to future positions on the faster timescale (∼125 ms) of the theta rhythm (“theta sequences”). During simulated offline rest, the network responded to transient noisy activation of random, sparse inputs by generating neuronal sequences that corresponded to forward, reverse, or mixed direction trajectories through the spatial environment.

Both online and offline sequence generation depended on structure in the strengths of excitatory synaptic connections such that pairs of neurons with overlapping spatial tuning were more strongly connected. In the online phase, different sparse subsets of excitatory neurons were activated at different spatial positions due to structure in the strengths of connections from spatially-tuned feedforward afferent inputs. The constraint that recurrent excitation must drive rhythmic synchronization in the theta band resulted in relatively strong recurrent connections. During each cycle of the theta rhythm, when the firing rates of the excitatory neurons were at their maximum, synaptic excitation from recurrent connections exceeded that from the feedforward afferents (Supplementary Figure S1E). This caused the sum of forward-moving feedforward inputs and symmetric mixed-direction feedback inputs to favor activation of cells encoding positions at or ahead of the current position. This generated forward-sweeping sequences that outpaced the speed of locomotion. However, at the opposite phase of the theta rhythm, when the firing rates of the excitatory cells reached their minimum, the non-rhythmic feedforward input became greater than recurrent excitation (Supplementary Figure S1E), causing theta sequences to reverse direction and relax back towards the current position encoded by the feedforward inputs.

In the offline phase, the feedforward inputs were not activated in a sequence, so momentum had to be entirely internally generated by the network. In this case, the particular subset of active feedforward inputs initially selected a sparse subset of excitatory neurons to begin to fire, which set a starting position for the replayed trajectory. Slight biases in the feedforward input could then influence whether the active ensemble of excitatory neurons next recruited neurons encoding spatial positions in either the forward or reverse direction. Once activity began moving in one direction, spike-rate adaptation facilitated continued sequence movement along that direction. However, depending on fluctuations in the feedforward inputs, sequences were also generated that included changes in direction. Interestingly, this process is akin to interpolation or smoothing – the recurrent connections within the network served to bridge large, discontinuous jumps in position encoded by the noisy feedforward inputs with smaller, more continuous steps. This produced offline sequences that were consistent with the topology of the spatial environment, but did not necessarily replay exact experienced trajectories. These findings are consistent with a recent report that neuronal sequences activated during hippocampal SWRs *in vivo* resembled Brownian motion, or a random walk through the sensory space, rather than precise replay of experience (Stella et al., 2019). This suggests that, rather than serving mainly to consolidate specific episodic memories of ordered sensory experiences, neuronal sequences during SWRs could also explore possible associations within the environment that had not been fully sampled during experience. Our modeling results showing that increased population representations of goal sites bias the content of offline memory replay also corroborate recent findings that previously rewarded locations are replayed more readily than immediate past or immediate future trajectories (Gillespie et al., 2021). Within this framework, synaptic plasticity during offline replay could modify connection strengths to increase the chance that a new path will be taken that is likely to lead to a desired outcome (Ólafsdóttir et al., 2015).

In summary, our modeling results identified a minimal set of elements sufficient to enable flexible and bidirectional memory replay in neuronal networks: spike rate adaptation, and recurrent connectivity between excitatory and inhibitory neuron populations with strengths and kinetics optimized for rhythmogenesis and sparse and selective stimulus representations. In previous models of neuronal sequence generation, additional network components were proposed to enable unidirectional sequences stored in memory to be reversed during offline recall, including neuromodulation (Gauy et al., 2018), excitability of neuronal dendrites (Gauy et al., 2018; Jahnke et al., 2015), coordinated plasticity at both excitatory and inhibitory synapses (Ramirez-Villegas et al., 2018), and functional specialization of diverse subpopulations of inhibitory interneurons (Cutsuridis and Hasselmo, 2011). While these mechanisms may regulate and enhance memory replay, our results suggest that they are not necessarily required.

This model also makes some experimentally-testable predictions. First, it implies that ion channel mutations that disrupt neuronal spike rate adaptation may also degrade neuronal sequence generation and memory consolidation (Peters et al., 2005). Secondly, while the direction and content of offline sequences may be largely controlled by internal dynamics and information stored in the synaptic weights within a recurrent neuronal circuit, the model network still required a small amount of random feedforward afferent input to evoke an offline population burst, suggesting that experimental manipulations of afferent projections to hippocampal area CA3 may alter the frequency or content of memory replay events (Chenani et al., 2019; Sasaki et al., 2018). Recent work has also begun to explore the advantages of generative replay for learning in artificial neural networks (Roscow et al., 2021). In addition to better understanding the biological mechanisms of memory consolidation and flexible planning of behavior, characterizing the minimal mechanisms of memory replay could facilitate the engineering of artificial systems that can refine their internal representations of the environment during periods of offline rest (Buzsáki, 1989).

## Acknowledgements

We are grateful to Kristopher Bouchard at LBNL for sharing large-scale computing resources provided by the National Energy Research Scientific Computing Center, a Department of Energy Office of Science User Facility (DE-AC02-05CH11231). This work was also made possible by computing allotments from NSF (XSEDE Comet, NCSA Blue Waters, and TACC Frontera) and supported by NIH BRAIN U19 grant NS104590 and NIMH grant R01MH121979.

## Materials and Methods

Simulations of a recurrent network of excitatory and inhibitory spiking neurons were executed using the python interface for the NEURON simulation software (Hines et al., 2009). Cell models were single-compartment integrate-and-fire neuronal cell models, as defined by Izhikevich (Izhikevich, 2007), and as implemented for the NEURON simulator by Lytton et al (Lytton et al., 2016). Previously calibrated cell models were replicated from those previous reports without modification – the “intrinsically bursting cell” model was used for excitatory neurons (E) with spike rate adaptation, the “regular spiking pyramidal cell” model was used for excitatory neurons without spike rate adaptation, and the “fast-spiking interneuron” model was used for inhibitory neurons (I) (Izhikevich, 2007; Lytton et al., 2016). Individual spikes in presynaptic neurons activated saturable conductance-based synapses with exponential rise and decay kinetics after a constant delay of 1 ms to emulate axonal conduction time (Carnevale and Hines, 2006). Excitatory synapses had a reversal potential of 0 mV (like AMPA-type glutamate receptors), and inhibitory synapses had a reversal potential of -80 mV (like GABA(A)-type receptors). In addition to the excitatory (E) and inhibitory (I) neuron populations, a population of feedforward afferent inputs (FF) provided a source of external excitatory synaptic drive to the model network.

The baseline weights of excitatory synapses onto E cells were sampled from a log-normal distribution (Almeida et al., 2009b; Buzsaki and Mizuseki, 2014), while the weights of excitatory synapses onto I cells, and all inhibitory synapses were sampled from a normal distribution (Grienberger et al., 2017). In addition to the random baseline synaptic weights assigned to excitatory synapses onto E cells, input strengths were increased by a variable additive factor that depended on the distance between the place fields of cells with overlapping spatial selectivity (Supplementary Figure S1B). The place field locations of the FF and E populations were assigned by distributing locations throughout the circular simulated track at equal intervals, and randomly assigning them to cells within each population. Random connectivity resulted in each E neuron receiving inputs from many FF and E neurons with heterogeneous selectivity, which produced substantial out-of-field excitation at all positions along the track.

For each of six types of connections between the three cell types (E <- FF, E <- E, E <- I, I <- FF, I <- E, I <- I), a number of parameters were varied and explored during optimization to identify model configurations that produced dynamics comparable to experimental observations. These parameters included: the mean and variance of the synaptic weight distribution for each connection type, the decay time constants of the synaptic conductances, the mean number of synapses made by one presynaptic cell onto one postsynaptic cell for each pair of cell types, and the maximum increase in synaptic weight due to shared selectivity, as mentioned above. Self-connections were not permitted.

Optimization was performed using a population-based iterative multi-objective algorithm. During each of 50 iterations, a population of 600 models with different parameters were simulated for one trial of simulated online run, and for 5 trials of simulated offline rest. During offline rest trials, a random subset of 25% of feedforward inputs were active with a mean rate of 12.5 Hz for an event duration of 160 ms (8 bins of 20 ms each). Different trials were implemented by using a distinct random number stream to sample unique spike times of the feedforward inputs from an inhomogeneous Poisson process. The following features of the network dynamics were evaluated for each model: average minimum and maximum firing rates of E cells during run, average mean firing rates of I cells during run, average fraction of active E and I cells during run, mean firing rates of E cells during rest, average fraction of active E cells during rest, and finally, features related to the frequency and power of theta and gamma band oscillations in E and I cells during run. These features were compared to target values to obtain a set of multiple objective error values. Models within a population were compared to each other and ranked with a non-dominated sorting procedure (Deb, 2011). Then, a new population of models was generated by making small perturbations to the parameter values of the most highly-ranked models from the previous iteration. This algorithm effectively identified model configurations that satisfied multiple objective criterion. Below, the final optimized parameter values (Table 1) and measured features of the network dynamics (Table 2) are compared for various model configurations discussed in this study:

**Table 2.** Features of model network dynamics.

| Feature | Target | Structured<br>E <- E<br>weights | Random<br>E <- E<br>weights | Shuffled<br>E <- E<br>weights | Suppressed<br>rhythmicity | No sparsity<br>or selectivity<br>constraints | No spike<br>rate<br>adaptation |
| --- | --- | --- | --- | --- | --- | --- | --- |
| E peak rate<br>(run) (Hz) | 20. | 17.20 | 19.96 | 10.42 | 20.07 | 19.73 | 7.30 |
| E min rate (run)<br>(Hz) | 0. | 0.24 | 0.25 | 0.27 | 0.25 | 12.37 | 0.39 |
| I mean rate<br>(run) (Hz) | 20. | 19.58 | 32.05 | 12.08 | 2.77 | 19.90 | 6.17 |
| E fraction<br>active (run) | 0.6 | 0.59 | 0.60 | 0.61 | 0.60 | 1.00 | 0.60 |
| I fraction active<br>(run) | 0.95 | 1.00 | 1.00 | 0.95 | 0.23 | 0.98 | 0.87 |
| E theta<br>amplitude<br>(run) | 0.5 | 0.78 | 0.37 | 0.63 | 0.15 | 0.77 | 1.29 |
| I theta<br>amplitude<br>(run) | 0.5 | 0.48 | 0.14 | 0.19 | 0.26 | 0.53 | 1.15 |
| E gamma<br>amplitude<br>(run) | 0.25 | 0.53 | 0.40 | 0.66 | 0.20 | 0.59 | 0.27 |
| I gamma<br>amplitude<br>(run) | 0.25 | 1.19 | 1.85 | 1.33 | 1.67 | 1.39 | 1.27 |
| E theta<br>frequency<br>(run) (Hz) | 7. | 7.09 | 7.45 | 7.64 | 11.27 | 7.09 | 6.73 |
| I theta<br>frequency<br>(run) (Hz) | 7. | 6.91 | 7.27 | 7.45 | 12.55 | 6.91 | 6.73 |
| E gamma<br>frequency<br>(run) (Hz) | 70. | 71.06 | 72.93 | 71.06 | 57.98 | 72.93 | 39.29 |
| I gamma<br>frequency<br>(run) (Hz) | 70. | 71.06 | 72.93 | 69.19 | 87.88 | 76.67 | 69.19 |
| E theta<br>frequency<br>tuning<br>index (run) | >5. | 6.40 | 6.30 | 7.00 | 0.00 | 29.05 | 150.67 |
| I theta<br>frequency<br>tuning<br>index (run) | >5. | 9.83 | 6.10 | 3.99 | 0.00 | 34.16 | 150.59 |
| E gamma<br>frequency<br>tuning<br>index (run) | >5. | 10.67 | 10.90 | 5.08 | -0.37 | 10.18 | -0.01 |
| I gamma<br>frequency | >5. | 16.16 | 4.14 | 31.65 | 0.36 | 6.02 | 3.43 |
| tuning index (run) |  |  |  |  |  |  |  |
| E fraction active (rest) | 0.25 | 0.37 | 0.20 | 0.53 | 0.36 | 1.00 | 0.40 |
| E mean rate (rest) (Hz) | 12.5 | 7.11 | 6.56 | 7.12 | 7.65 | 17.77 | 6.97 |

In the above table, theta and gamma amplitudes were quantified as follows: average population firing rates were band-pass filtered, and the envelopes of the filtered traces were computed from the Hilbert transformation. Then power was expressed as a ratio of the average envelope amplitude to the average population firing rate. To quantify theta and gamma frequency, bandpass filtered traces were subject to frequency decomposition, and the frequency corresponding to the centroid or center-of-mass of the power spectral density distribution was taken as the dominant frequency within the band. The area of the power spectral density distribution was also used to compute a “frequency tuning index” which quantified how concentrated the power distribution was around the centroid frequency. This metric was akin to a signal-to-noise ratio in the frequency domain instead of the time domain, and was computed as follows:

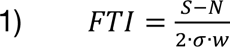

where 𝑆 is the average power at frequencies within the center of mass quartile containing the centroid frequency (signal), 𝑁 is the average power at frequencies in the extreme high and low quartiles outside the center of mass quartile (noise), 𝜎 is the standard deviation of the power distribution, and 𝑤 is the half-width of the power distribution in the frequency domain normalized to the width of the bandpass filter. This metric has values near zero when power is distributed uniformly within the filter band, and values larger than one when power is concentrated around the centroid frequency.

Following parameter optimization, each variant of the network was evaluated by simulating 5 trials of online run, and 1000 trials of offline rest for each of 5 independent network instances. For a given set of model parameters, independent instances of each network variant were constructed by using distinct random number streams to assign place field locations, input spike times, synaptic connections, and synaptic weights for all cells in the network.

Bayesian decoding of spatial position from spike times recorded during a single trial (Figures 2, 3, 5 and 6, and Supplementary Figures S3 – S6) was performed using the procedure described in (Davidson et al., 2009). The spatial firing rates of all cells were averaged across 5 trials of simulated online run to compute the spiking probabilities of each neuron in 20 ms bins. Then, spiking data was taken from either a held-out set of 5 trials of simulated run (Figure 2), or offline rest trials (Figures 3, 5 and 6, and Supplementary Figures S3 – S6). The numbers of spikes emitted by each cell in 20 ms bins were used to determine a likelihood distribution over spatial positions. The position with maximum likelihood was used as the decoded position estimate for each temporal bin. In Figure 2A, decoded positions of E and I cells sweep smoothly from past to future positions, and then relax back to the current position on the timescale of the ongoing theta rhythm. To quantify this form of online neuronal sequence generation, a theta sequence score (Figures 2E and 4H) was computed as follows: decoded position error was first bandpass filtered in the theta band (4 – 10 Hz). Then the contribution of this oscillation to the total variance in decoded position error was calculated as the square of the correlation (R^2^) between the original mean-subtracted error signal and the theta filtered signal. In Figures 3F, 3M, 4I, 5G and 6G, and Supplementary Figures S3H, S4F, S5K, S6K and S8E, offline sequences were categorized as consistent with a continuous trajectory through space if they met the following criterion: 1) at least one cell in a population must emit at least one spike in each temporal bin, 2) the change in decoded position between any two adjacent bins must not exceed 35% of the track length, 3) the total path length of the decoded trajectory must not exceed 100% of the track length, and 4) the net speed of the trajectory (absolute value of net change in position divided by the 160 ms offline event duration) must exceed 50% of the run speed of 0.33 track lengths / sec used during simulation of online exploration.

In Figure 4 and Supplementary Figure S8, the diversity and degeneracy of various model configurations was explored as follows: for each model configuration, 30,000 models were evaluated during parameter optimization, and the model with the lowest multi-objective error score was considered the “best” model. The remaining models were sorted by their Euclidean distance from the “best” model in the space of model parameters. This resulted in an error landscape (e.g. Figure 4A) in which models with similar parameters resulted in similar multi-objective error scores. We then identified models located at local minima in this error landscape, which as a group comprised models that were distant from each other in parameter space, but similar to each other in terms of overall multi-objective error. We further enforced that selected models had to be a minimum distance of 0.15 from each other in parameter space, and selected 5 such models with the lowest error score to be included in a “Marder group” of models for further analysis. For each alternative network model configuration (i.e. network models with and without structured recurrent excitatory connections), each of 5 “Marder group” model variants with different parameters were evaluated for offline sequence generation by simulating 1000 trials for each of 5 independent network instances.

In box and whisker plots in Figures 2D, 2E, 3F, 3M, 4D – 4I, 5G and 6G, and Supplementary Figures S1C, S1D, S3H, S4F, S5K, S6K and S8A – S8E, center lines indicate median, boxes span the first and third quartile of the data, and whiskers extend to 1.5 times the inter-quartile range.

### Data and Code Availability

All code necessary to reproduce the data and analysis presented in this work are available here:

Network simulation and analysis code: https://github.com/neurosutras/optimize_simple_network

Network optimization code: https://github.com/neurosutras/nested

---

**Supplementary Figure S1.**
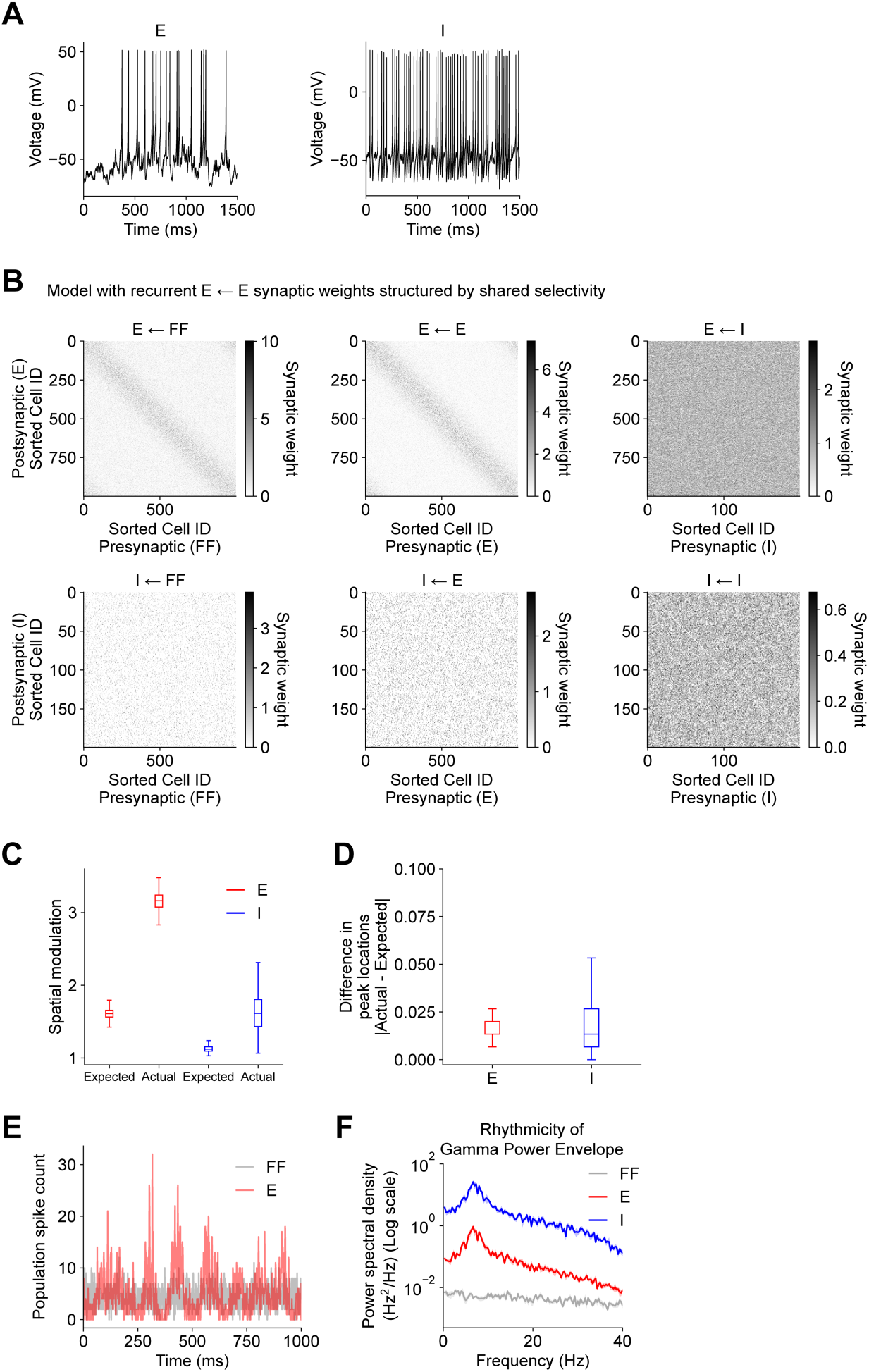
Related to Figures 1, 2 and 3. Structure and online dynamics of network model with excitatory synaptic connectivity structured by shared stimulus selectivity. (**A**) Intracellular voltage recordings from example E and I cells during simulated “online” spatial exploration. (**B**) The strengths of synaptic connections within the network model shown in Figure 1 are indicated by grayscale intensity. Cells in each population are sorted by the location of maximum average spatial firing rate. Excitatory connections from FF and E cells onto E cells are increased in strength for pairs of cells with overlapping spatial selectivity. (**C**) Spatial modulation for each cell is computed as a ratio of maximum to mean activity. The degree of spatial modulation expected from a linear weighted sum of excitatory inputs is larger for E cells than I cells due to the structure of the weight distributions shown in (B). However, the actual spatial modulation measured from the firing rate outputs of each cell is larger than expected in both E and I cells due to suppression of background activity by synaptic inhibition (one-sided paired t-tests: Expected vs. Actual: E, p<0.00001; I, p<0.00001). (**D**) The distance between the expected location of maximum firing, and the actual location is quantified as a fraction of the circular track (one-sample t-tests: E, p<0.00001; I, p=0.00005). (**E**) Traces depict average population firing rates of E and FF cells during an example trial for one instance of the network. E cells and FF cells dominate at different phases of the population theta oscillation. (**F**) In Figure 1G (bottom row), the amplitudes of the gamma-filtered (purple) population firing rates for E and I populations vary in time. Here, the amplitudes or envelopes of the gamma-filtered population firing rates were subject to frequency decomposition. The resulting frequency distributions show peaks in the theta band (one-sided paired t-tests: 4 Hz – 10 Hz frequency band: E vs. FF, p=0.00002; I vs. FF, p=0.00002). Mean (solid) ± SEM (shading) were computed across 5 independent network instances. In (C) and (D), box and whisker plots depict data across cells for one example instance of the network (see Materials and Methods). Statistics were computed across 5 independent instances of the network. p-values reflect FDR correction for multiple comparisons.

**Supplementary Figure S2.**
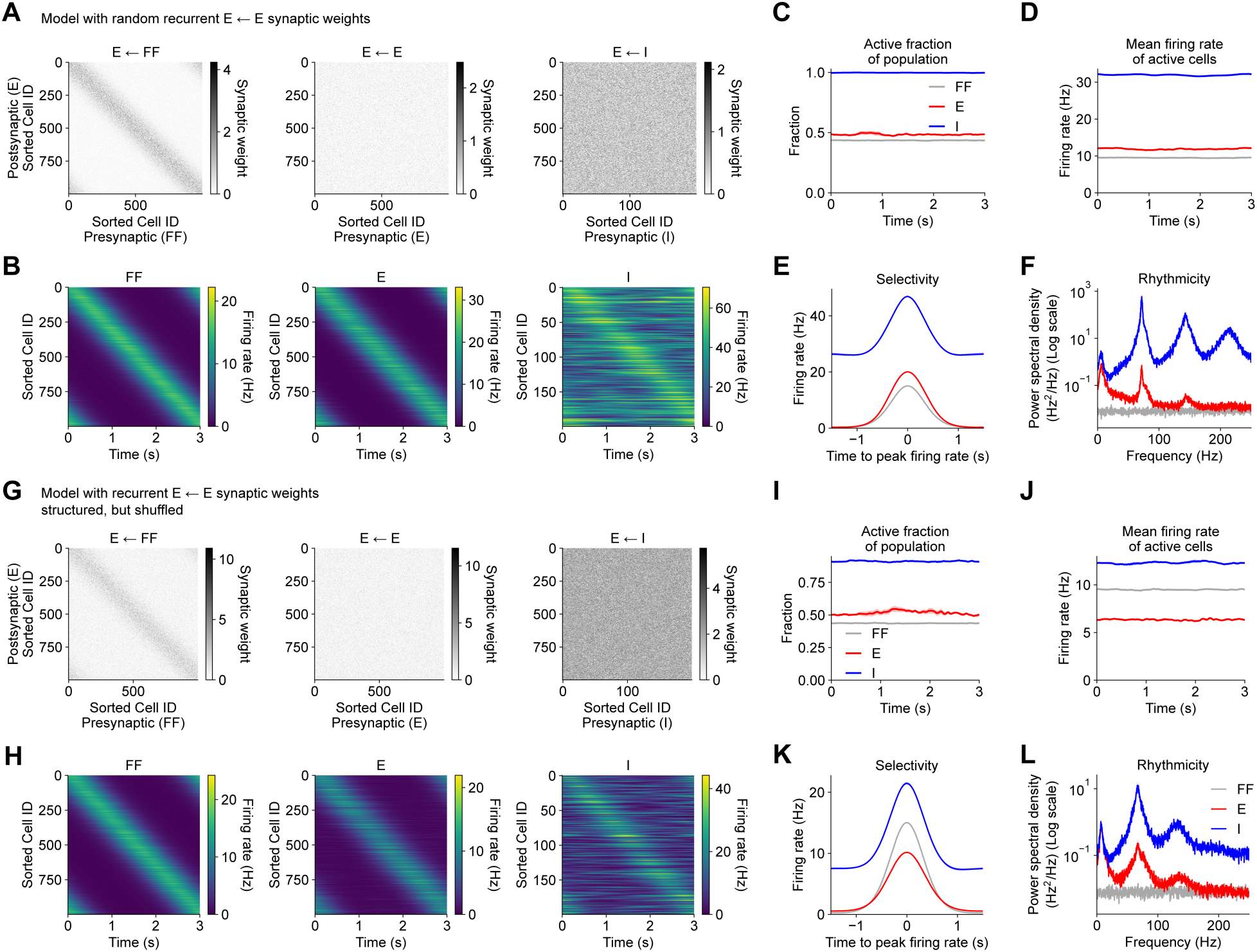
Related to Figures 1, 2 and 3. Structure and online dynamics of alternative network models with random recurrent excitatory connectivity. (**A**) Same as Supplementary Figure S1B for an alternative network model with random E ← E weights. (**B** – **F**) Same as Figures 1C – 1F and 1H for alternative network model. (**F**) Rhythmicity (one-sided paired t-tests: theta: E vs. FF, p=0.00003; I vs. FF, p<0.00001; gamma: E vs. FF, p<0.00001; I vs. FF, p<0.00001; two-sided t- tests vs. data from model with structured E ← E weights in Figure 1H: theta: E, p<0.00001; I, p<0.00001; gamma: E, p<0.00001; I, p<0.00001). (**G** – **L**) Same as (A – F) for an alternative network model with structured, but shuffled E ← E weights. (**H**) Rhythmicity (one-sided paired t-tests: theta: E vs. FF, p=0.00009; I vs. FF, p=0.00003; gamma: E vs. FF, p=0.00002; I vs. FF, p<0.00001; two-sided t-tests vs. data from model with structured E ← E weights in Figure 1H: theta: E, p<0.00001; I, p<0.00001; gamma: E, p=0.00001; I, p<0.00001). In (C – F) and (I – L), data were first averaged across 5 trials per network instance. Mean (solid) ± SEM (shading) were computed across 5 independent network instances. p-values reflect FDR correction for multiple comparisons.

**Supplementary Figure S3.**
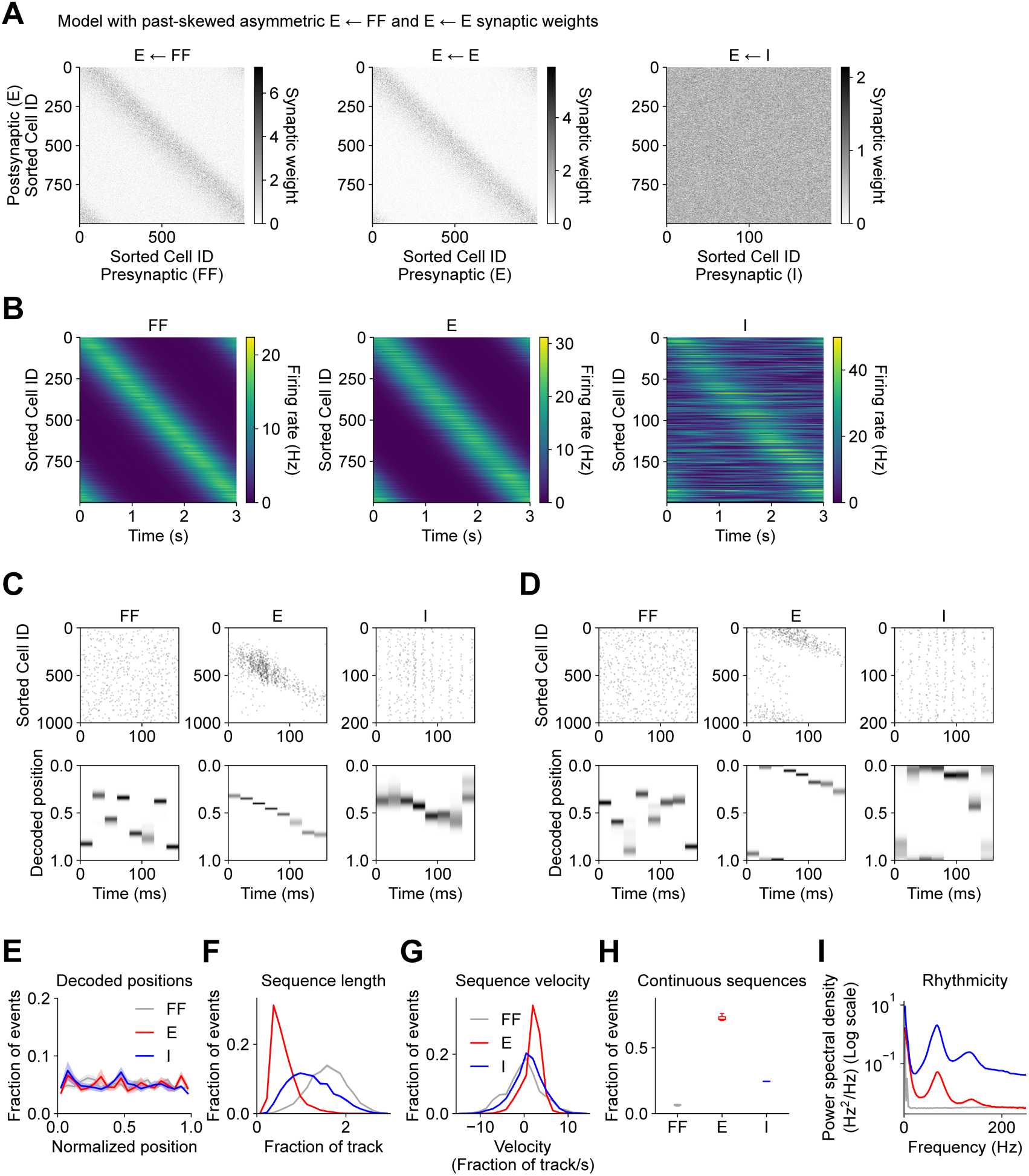
Related to Figures 1 and 3. Offline replay is unidirectional in an alternative network model with asymmetric synaptic connectivity. (**A**) Same as Supplementary Figure S1B for an alternative network model with E ← E and E ← FF synaptic weights biased such that neurons encoding the current position of the animal during run preferentially activate neurons encoding future positions. This biases offline replay towards forward-moving unidirectional sequences. (**B**) Same as Figure 1C for alternative network model. (**C** – **I**) Same as Figures 3A – 3G for alternative network model. (**C**) and (**D**) correspond to two example trials from one example network instance. (**E**) Decoded positions (two-sided K-S tests: E vs. FF, p=0.99974; I vs. FF, p=0.99974; two-sided K-S tests vs. data from model with structured E ← E weights in Figure 3C: E, p=0.99974; I, p=0.99974). (**F**) Offline sequence length (two-sided K-S tests: E vs. FF, p<0.00001; I vs. FF, p<0.00001; two-sided K-S tests vs. data from model with structured E ← E weights in Figure 3D: E, p<0.00001; I, p=0.31278). (**G**) Offline sequence velocity (two-sided K-S tests: E vs. FF, p<0.00001; I vs. FF, p=0.00026; two-sided K-S tests vs. data from model with structured E ← E weights in Figure 3E: E, p<0.00001; I, p=0.00026). (**H**) Fraction of events that met criterion for sequences consistent with continuous spatial trajectories (see Materials and Methods) (one-sided paired t-tests: E vs. FF, p<0.00001, I vs. FF, p=0.00003; two-sided t-tests vs. data from model with structured E ← E weights in Figure 3F: E, p<0.00001; I, p=0.00112). (**I**) Offline high-frequency rhythmicity (one-sided paired t-tests: 75 Hz – 300 Hz frequency band: E vs. FF, p=0.00100; I vs. FF, p=0.00001; two-sided t-tests vs. data from model with structured E ← E weights in Figure 3G: E, p=0.00001; I, p=0.02509). In (E – G) and (I), mean (solid) ± SEM (shading) were computed across 5 independent instances of each network. In (H), box and whisker plots depict data from 5 independent instances of each network model (see Materials and Methods). p-values reflect FDR correction for multiple comparisons.

**Supplementary Figure S4.**
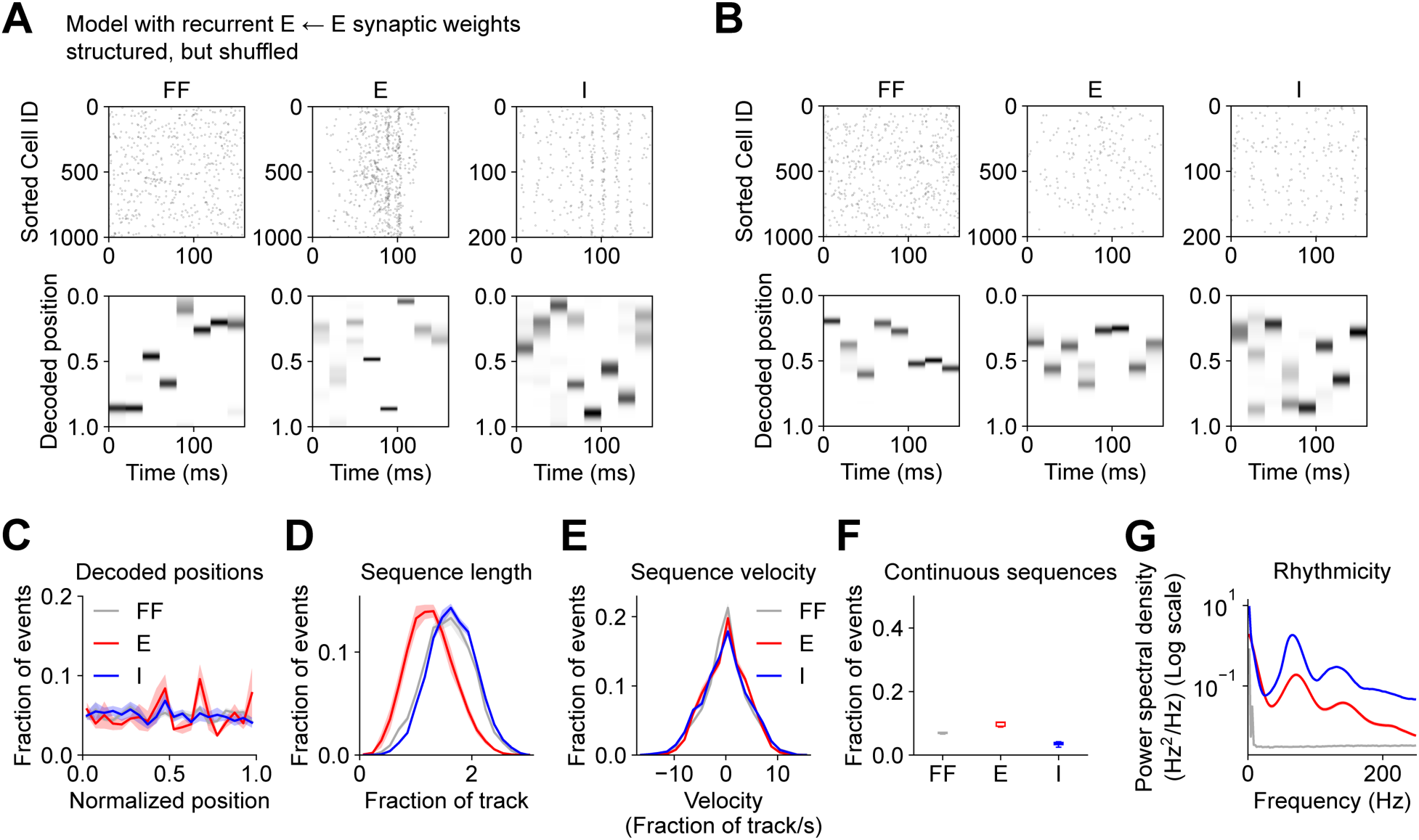
Related to Figure 3. Offline replay is disrupted in an alternative network model with structured, but shuffled recurrent excitatory connectivity. (**A** – **G**) Same as Figures 3A – 3G for an alternative network model with structured, but shuffled E ← E weights. (**A**) and (**B**) correspond to two example trials from one example network instance. (**C**) Decoded positions (two-sided K-S tests: E vs. FF, p=0.99974; I vs. FF, p=0.99974; two-sided K-S tests vs. data from model with structured E ← E weights in Figure 3C: E, p=0.99974; I, p=0.99974). (**D**) Offline sequence length (two-sided K-S tests: E vs. FF, p<0.00001; I vs. FF, p=0.11015; two-sided K-S tests vs. data from model with structured E ← E weights in Figure 3D: E, p<0.00001; I, p<0.00001). (**E**) Offline sequence velocity (two-sided K-S tests: E vs. FF, p=0.96717; I vs. FF, p=0.73515; two-sided K-S tests vs. data from model with structured E ← E weights in Figure 3E: E, p=0.00272; I, p=0.24955). (**F**) Fraction of events that met criterion for sequences consistent with continuous spatial trajectories (see Materials and Methods) (one-sided paired t-tests: E vs. FF, p=0.06713, I vs. FF, p=0.99967; two-sided t-tests vs. data from model with structured E ← E weights in Figure 3F: E, p<0.00001; I, p<0.00001). (**G**) Offline high-frequency rhythmicity (one-sided paired t-tests: 75 Hz – 300 Hz frequency band: E vs. FF, p=0.00092; I vs. FF, p=0.00004; two-sided t-tests vs. data from model with structured E ← E weights in Figure 3G: E, p=0.00006; I, p=0.00059). In (C – E) and (G), mean (solid) ± SEM (shading) were computed across 5 independent instances of each network. In (F), box and whisker plots depict data from 5 independent instances of each network model (see Materials and Methods). p-values reflect FDR correction for multiple comparisons.

**Supplementary Figure S5.**
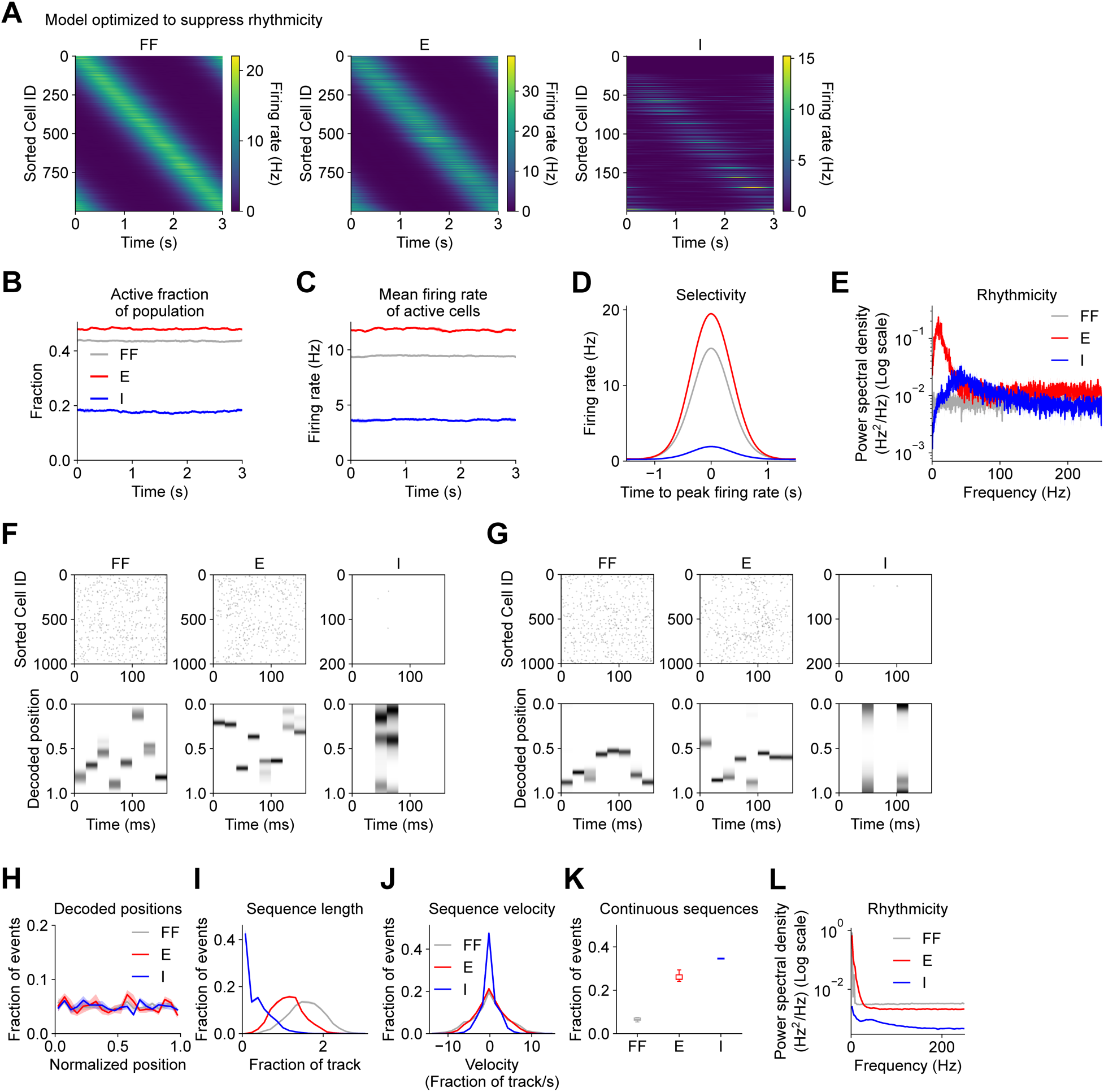
Related to Figures 1 and 3. Offline replay is disrupted in an alternative network model optimized to suppress rhythmicity. (**A** – **E**) Same as Figures 1C – 1F and 1H for alternative network model optimized to suppress theta and gamma rhythmicity in E and I cells. (**E**) Rhythmicity (one-sided paired t-tests: theta: E vs. FF, p<0.00001; I vs. FF, p=0.99754; gamma: E vs. FF, p<0.00001; I vs. FF, p=0.00003; two-sided t-tests vs. data from model with structured E ← E weights in Figure 1H: theta: E, p<0.00001; I, p<0.00001; gamma: E, p<0.00001; I, p<0.00001). (**F** – **L**) Same as Figures 3A – 3G for alternative network model. (**F**) and (**G**) correspond to two example trials from one example network instance. (**H**) Decoded positions (two-sided K-S tests: E vs. FF, p=0.99974; I vs. FF, p=0.99974; two-sided K-S tests vs. data from model with structured E ← E weights in Figure 3C: E, p=0.99974; I, p=0.99974). (**I**) Offline sequence length (two-sided K-S tests: E vs. FF, p<0.00001; I vs. FF, p<0.00001; two-sided K-S tests vs. data from model with structured E ← E weights in Figure 3D: E, p<0.00001; I, p<0.00001). (**J**) Offline sequence velocity (two-sided K-S tests: E vs. FF, p=0.73515; I vs. FF, p=0.00001; two-sided K-S tests vs. data from model with structured E ← E weights in Figure 3E: E, p=0.09901; I, p=0.00004). (**K**) Fraction of events that met criterion for sequences consistent with continuous spatial trajectories (see Materials and Methods) (one-sided paired t-tests: E vs. FF, p=0.00004, I vs. FF, p=0.00002; two-sided t-tests vs. data from model with structured E ← E weights in Figure 3F: E, p<0.00001; I, p=0.00001). (**L**) Offline high-frequency rhythmicity (one-sided paired t-tests: 75 Hz – 300 Hz frequency band: E vs. FF, p=1; I vs. FF, p=1; two-sided t-tests vs. data from model with structured E ← E weights in Figure 3G: E, p=0.00001; I, p<0.00001). In (B – E), (H – J) and (L), mean (solid) ± SEM (shading) were computed across 5 independent instances of each network. In (K), box and whisker plots depict data from 5 independent instances of each network model (see Materials and Methods). p-values reflect FDR correction for multiple comparisons.

**Supplementary Figure S6.**
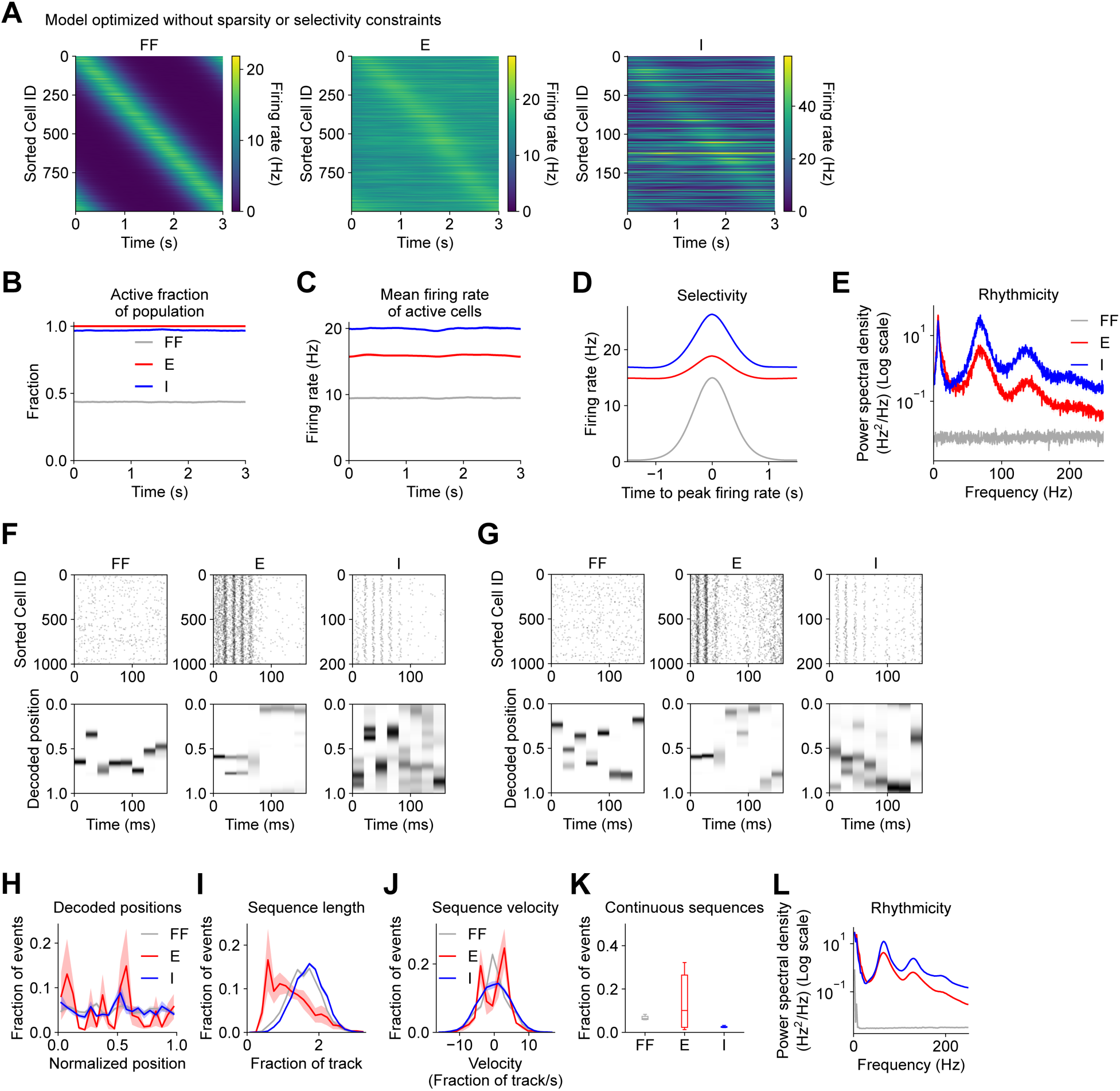
Related to Figures 1 and 3. Offline replay is disrupted in an alternative network model optimized without sparsity or selectivity constraints. (**A** – **E**) Same as Figures 1C – 1F and 1H for alternative network model optimized without sparsity or selectivity constraints in E cells. (**E**) Rhythmicity (one-sided paired t-tests: theta: E vs. FF, p<0.00001; I vs. FF, p<0.00001; gamma: E vs. FF, p<0.00001; I vs. FF, p<0.00001; two-sided t-tests vs. data from model with structured E ← E weights in Figure 1H: theta: E, p<0.00001; I, p<0.00001; gamma: E, p<0.00001; I, p<0.00001). (**F** – **L**) Same as Figures 3A – 3G for alternative network model. (**F**) and (**G**) correspond to two example trials from one example network instance. (**H**) Decoded positions (two-sided K-S tests: E vs. FF, p=0.00019; I vs. FF, p=0.99974; two-sided K-S tests vs. data from model with structured E ← E weights in Figure 3C: E, p=0.00010; I, p=0.99974). (**I**) Offline sequence length (two-sided K-S tests: E vs. FF, p<0.00001; I vs. FF, p=0.00030; two-sided K-S tests vs. data from model with structured E ← E weights in Figure 3D: E, p<0.00001; I, p<0.00001). (**J**) Offline sequence velocity (two-sided K-S tests: E vs. FF, p=0.00015; I vs. FF, p=0.31270; two-sided K-S tests vs. data from model with structured E ← E weights in Figure 3E: E, p=0.00093; I, p=0.06213). (**K**) Fraction of events that met criterion for sequences consistent with continuous spatial trajectories (see Materials and Methods) (one-sided paired t-tests: E vs. FF, p=0.14499, I vs. FF, p=0.99967; two-sided t-tests vs. data from model with structured E ← E weights in Figure 3F: E, p=0.00027; I, p<0.00001). (**L**) Offline high-frequency rhythmicity (one-sided paired t-tests: 75 Hz – 300 Hz frequency band: E vs. FF, p<0.00001; I vs. FF, p<0.00001; two-sided t-tests vs. data from model with structured E ← E weights in Figure 3G: E, p<0.00001; I, p<0.00001). In (B – E), (H – J) and (L), mean (solid) ± SEM (shading) were computed across 5 independent instances of each network. In (K), box and whisker plots depict data from 5 independent instances of each network model (see Materials and Methods). p-values reflect FDR correction for multiple comparisons.

**Supplementary Figure S7.**
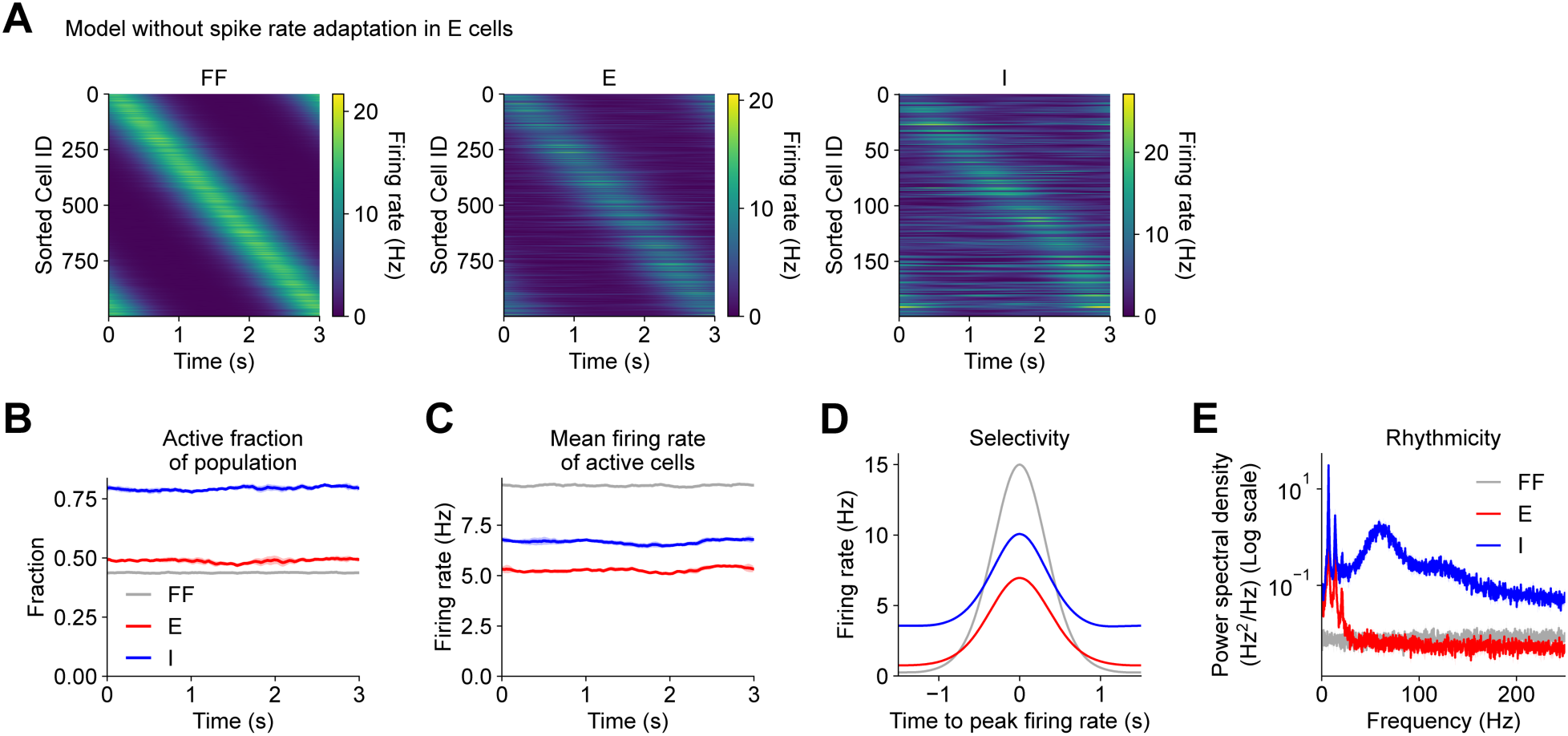
Related to Figures 1 and 5. Online dynamics in network model without spike rate adaptation. (**A** – **E**) Same as Figures 1C – 1F and 1H for alternative network model without spike rate adaptation in E cells. (**E**) Rhythmicity (one-sided paired t-tests: theta: E vs. FF, p=0.00016; I vs. FF, p=0.00007; gamma: E vs. FF, p=0.98658; I vs. FF, p=0.00002; two-sided t-tests vs. data from model with structured E ← E weights in Figure 1H: theta: E, p=0.03333; I, p<0.00001; gamma: E, p<0.00001; I, p<0.00001). In (B – E), mean (solid) ± SEM (shading) were computed across 5 independent instances of each network. p-values reflect FDR correction for multiple comparisons.

**Supplementary Figure S8.**
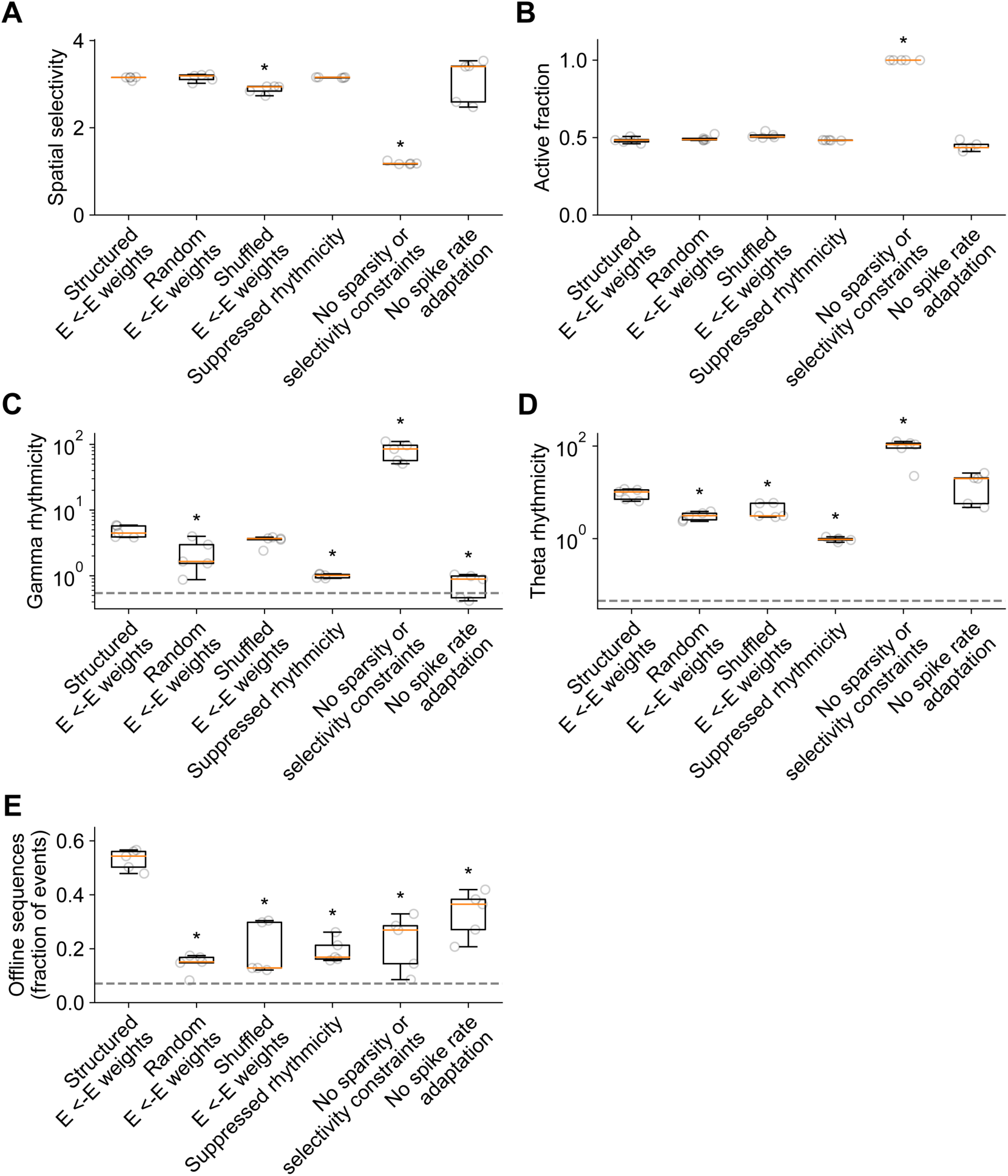
Related to Figures 1, 3, 4, and 5. Exploration of model parameter diversity and degeneracy in additional alternative network models. To explore model diversity and degeneracy, for various network model configurations, a subset of model variants termed a “Marder group” were selected based on their large distance from each other in the space of parameters, but their similar performance with respect to multiple optimization objectives (see Figure 4 and Materials and Methods). Features of the simulated network dynamics produced by distinct model variants within a “Marder group” are compared across network model configurations. (**A**) Spatial selectivity of the excitatory neuron population during simulated “online exploration” is computed as a ratio of maximum to mean activity (two-sided t-tests vs. data from model with structured E ← E weights: Random E ← E weights: p=1; Shuffled E ← E weights: p=0.00224; No rhythmicity constraints: p=1; No sparsity or selectivity constraints: p<0.00001; No spike rate adaptation: p=1). (**B**) The fraction of the excitatory neuron population that is synchronously active during simulated “online exploration” is shown (two-sided t-tests vs. data from model with structured E ← E weights: Random E ← E weights: p=1; Shuffled E ← E weights: p=0.12079; No rhythmicity constraints: p=1; No sparsity or selectivity constraints: p<0.00001; No spike rate adaptation: p=0.19538). (**C**) Gamma rhythmicity of the excitatory neuron population is computed as the integrated power spectral density in the gamma frequency band (two-sided t-tests vs. data from model with structured E ← E weights: Random E ← E weights: p=0.03552; Shuffled E ← E weights: p=0.16364; No rhythmicity constraints: p=0.00014; No sparsity or selectivity constraints: p=0.00091; No spike rate adaptation: p=0.00012). (**D**) Theta rhythmicity of the excitatory neuron population is computed as the integrated power spectral density in the theta frequency band (two-sided t-tests vs. data from model with structured E ← E weights: Random E ← E weights: p=0.00272; Shuffled E ← E weights: p=0.01944; No rhythmicity constraints: p=0.00030; No sparsity or selectivity constraints: p=0.00984; No spike rate adaptation: p=1). (**E**) Fraction of events during simulated “offline rest” that met criterion for sequences consistent with continuous spatial trajectories (see Figure 3, Supplementary Figure S3, and Materials and Methods) (two-sided t-tests vs. data from model with structured E ← E weights: Random E ← E weights: p<0.00001; Shuffled E ← E weights: p=0.00044; No rhythmicity constraints: p=<0.00001; No sparsity or selectivity constraints: p=0.00120; No spike rate adaptation: p=0.00751). In (A – E), for each network model configuration, box and whisker plots depict 5 distinct “Marder group” model variants with different parameters (see Materials and Methods). Data for each model variant (grey circles) were first averaged across 5 independent instances of that model variant. For each network model configuration, In (C – E), grey dashed lines indicate value for the feedforward input to the network for reference. p-values reflect Bonferroni correction for multiple comparisons.

**Supplementary Figure S9.**
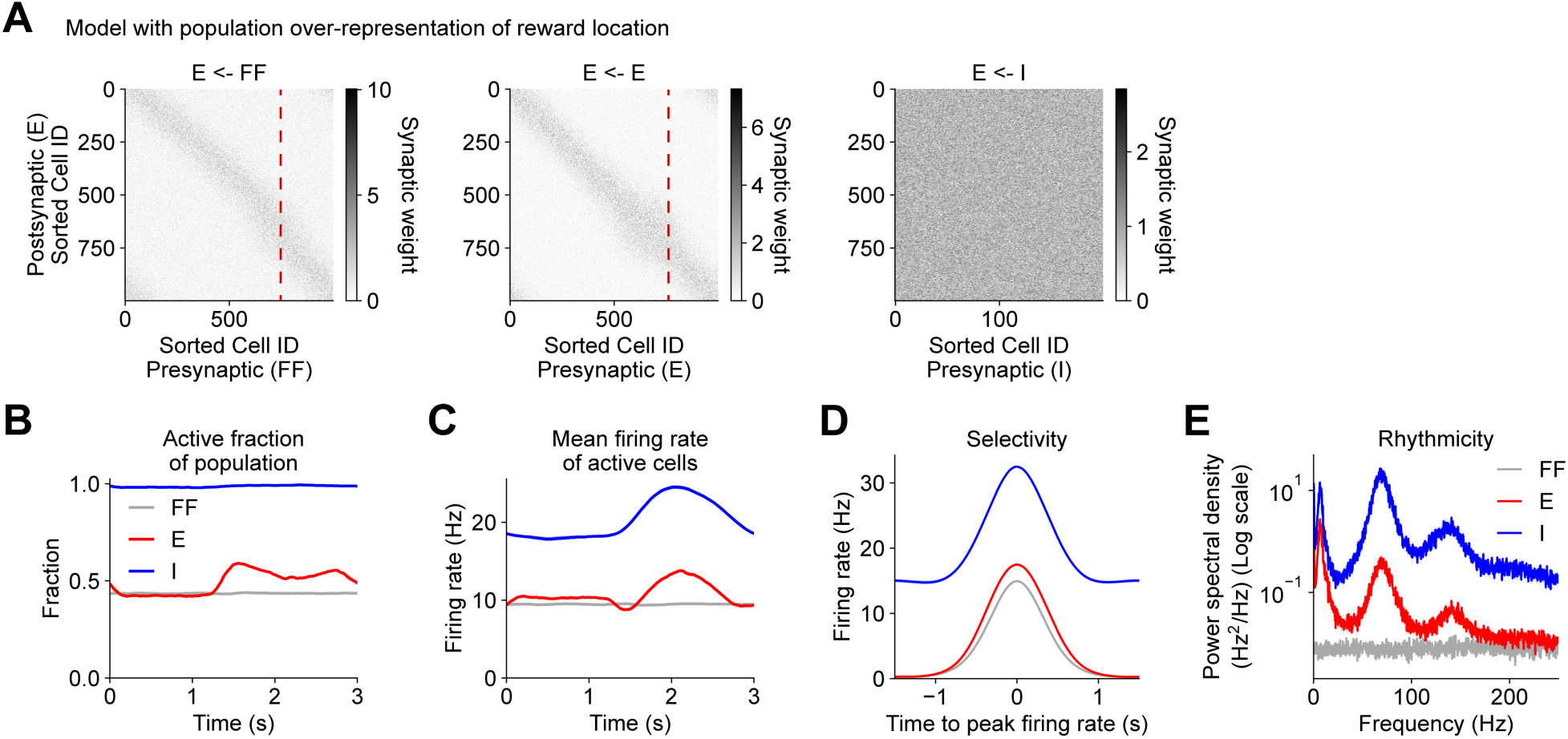
Related to Figures 1 and 6. Online activity in network model with population over-representation of reward. (**A**) Same as Supplementary Figure S1B for an alternative network model with population-level over-representation of reward location in E cells. Simulated reward site is marked with red dashed line. (**B** – **E**) Same as Figures 1D – 1F and 1H for alternative network model. (**E**) Rhythmicity (one-sided paired t-tests: theta: E vs. FF, p<0.00001; I vs. FF, p<0.00001; gamma: E vs. FF, p=0.00003; I vs. FF, p<0.00001; two-sided t-tests vs. data in Figure 1H: theta: E, p=0.00440; I, p=0.16525; gamma: E, p=0.00043; I, p=0.01695). In (B – E), mean (solid) ± SEM (shading) were computed across 5 independent instances of each network. p-values reflect FDR correction for multiple comparisons.

## References

Almeida, L. de, Idiart, M., and Lisman, J.E. (2007). Memory retrieval time and memory capacity of the CA3 network: role of gamma frequency oscillations. Learn Memory 14, 795–806. https://doi.org/10.1101/lm.730207.

Almeida, L. de, Idiart, M., and Lisman, J.E. (2009a). A second function of gamma frequency oscillations: an E%-max winner-take-all mechanism selects which cells fire. J Neurosci 29, 7497–7503. https://doi.org/10.1523/jneurosci.6044-08.2009.

Almeida, L. de, Idiart, M., and Lisman, J.E. (2009b). The input-output transformation of the hippocampal granule cells: from grid cells to place fields. J Neurosci 29, 7504– 7512. https://doi.org/10.1523/jneurosci.6048-08.2009.

Amit, D.J., and Brunel, N. (1997). Dynamics of a recurrent network of spiking neurons before and following learning. Netw Comput Neural Syst 8, 373–404. https://doi.org/10.1088/0954-898x_8_4_003.

Arkhipov, A., Gouwens, N.W., Billeh, Y.N., Gratiy, S., Iyer, R., Wei, Z., Xu, Z., Abbasi-Asl, R., Berg, J., Buice, M., et al. (2018). Visual physiology of the layer 4 cortical circuit in silico. Plos Comput Biol 14, e1006535. https://doi.org/10.1371/journal.pcbi.1006535.

Bezaire, M.J., Raikov, I., Burk, K., Vyas, D., and Soltesz, I. (2016). Interneuronal mechanisms of hippocampal theta oscillation in a full-scale model of the rodent CA1 circuit. Elife 5, e18566. https://doi.org/10.7554/elife.18566.

Bittner, K.C., Grienberger, C., Vaidya, S.P., Milstein, A.D., Macklin, J.J., Suh, J., Tonegawa, S., and Magee, J.C. (2015). Conjunctive input processing drives feature selectivity in hippocampal CA1 neurons. Nat Neurosci 18, 1133–1142. https://doi.org/10.1038/nn.4062.

Bittner, K.C., Milstein, A.D., Grienberger, C., Romani, S., and Magee, J.C. (2017). Behavioral time scale synaptic plasticity underlies CA1 place fields. Science 357, 1033–1036. https://doi.org/10.1126/science.aan3846.

Brunel, N. (2016). Is cortical connectivity optimized for storing information? Nat Neurosci 19, 749–755. https://doi.org/10.1038/nn.4286.

Brunel, N., and Trullier, O. (1998). Plasticity of directional place fields in a model of rodent CA3. Hippocampus 8, 651–665. https://doi.org/10.1002/(sici)1098-1063(1998)8:6<651::aid-hipo8>3.0.co;2-l.

Brunel, N., and Wang, X.-J. (2003). What Determines the Frequency of Fast Network Oscillations With Irregular Neural Discharges? I. Synaptic Dynamics and Excitation-Inhibition Balance. J Neurophysiol 90, 415–430. https://doi.org/10.1152/jn.01095.2002.

Buzsáki, G. (1989). Two-stage model of memory trace formation: A role for “noisy” brain states. Neuroscience 31, 551–570. https://doi.org/10.1016/0306-4522(89)90423-5.

Buzsaki, G., and Mizuseki, K. (2014). The log-dynamic brain: how skewed distributions affect network operations. Nat Rev Neurosci 15, 264–278. https://doi.org/10.1038/nrn3687.

Buzsáki, G., McKenzie, S., and Davachi, L. (2021). Neurophysiology of Remembering. Annu Rev Psychol 73, 1–29. https://doi.org/10.1146/annurev-psych-021721-110002.

Carnevale, N.T., and Hines, M.L. (2006). The NEURON Book (Cambridge University Press).

Chadwick, A., Rossum, M.C. van, and Nolan, M.F. (2015). Independent theta phase coding accounts for CA1 population sequences and enables flexible remapping. Elife 4, e03542. https://doi.org/10.7554/elife.03542.

Chadwick, A., Rossum, M.C. van, and Nolan, M.F. (2016). Flexible theta sequence compression mediated via phase precessing interneurons. Elife 5, e20349. https://doi.org/10.7554/elife.20349.

Chenani, A., Sabariego, M., Schlesiger, M.I., Leutgeb, J.K., Leutgeb, S., and Leibold, C. (2019). Hippocampal CA1 replay becomes less prominent but more rigid without inputs from medial entorhinal cortex. Nat Commun 10, 1341. https://doi.org/10.1038/s41467-019-09280-0.

Colgin, L.L. (2016). Rhythms of the hippocampal network. Nat Rev Neurosci 17, 239– 249. https://doi.org/10.1038/nrn.2016.21.

Csicsvari, J., Hirase, H., Mamiya, A., and Buzsaki, G. (2000). Ensemble patterns of hippocampal CA3-CA1 neurons during sharp wave-associated population events. Neuron 28, 585–594. .

Csicsvari, J., Jamieson, B., Wise, K.D., and Buzsaki, G. (2003). Mechanisms of gamma oscillations in the hippocampus of the behaving rat. Neuron 37, 311–322. .

Cutsuridis, V., and Hasselmo, M. (2011). Spatial Memory Sequence Encoding and Replay During Modeled Theta and Ripple Oscillations. Cogn Comput 3, 554–574. https://doi.org/10.1007/s12559-011-9114-3.

Davidson, T.J., Kloosterman, F., and Wilson, M.A. (2009). Hippocampal replay of extended experience. Neuron 63, 497–507. https://doi.org/10.1016/j.neuron.2009.07.027.

Deb, K. (2011). Multi-objective Evolutionary Optimisation for Product Design and Manufacturing. 3–34. https://doi.org/10.1007/978-0-85729-652-8_1.

Dorkenwald, S., Turner, N.L., Macrina, T., Lee, K., Lu, R., Wu, J., Bodor, A.L., Bleckert, A.A., Brittain, D., Kemnitz, N., et al. (2019). Binary and analog variation of synapses between cortical pyramidal neurons. Biorxiv 2019.12.29.890319. https://doi.org/10.1101/2019.12.29.890319.

Dragoi, G., and Tonegawa, S. (2013). Distinct preplay of multiple novel spatial experiences in the rat. Proc National Acad Sci 110, 9100–9105. https://doi.org/10.1073/pnas.1306031110.

Drieu, C., and Zugaro, M. (2019). Hippocampal Sequences During Exploration: Mechanisms and Functions. Front Cell Neurosci 13, 232. https://doi.org/10.3389/fncel.2019.00232.

Duigou, C.L., Simonnet, J., Teleñczuk, M.T., Fricker, D., and Miles, R. (2014). Recurrent synapses and circuits in the CA3 region of the hippocampus: an associative network. Front Cell Neurosci 7, 262. https://doi.org/10.3389/fncel.2013.00262.

Ego-Stengel, V., and Wilson, M.A. (2007). Spatial selectivity and theta phase precession in CA1 interneurons. Hippocampus 17, 161–174. https://doi.org/10.1002/hipo.20253.

Ermentrout, B. (1992). Complex dynamics in winner-take-all neural nets with slow inhibition. Neural Networks 5, 415–431. https://doi.org/10.1016/0893-6080(92)90004-3.

Farooq, U., and Dragoi, G. (2019). Emergence of preconfigured and plastic time-compressed sequences in early postnatal development.

Fernández-Ruiz, A., Oliva, A., Oliveira, E.F. de, Rocha-Almeida, F., Tingley, D., and Buzsáki, G. (2019). Long-duration hippocampal sharp wave ripples improve memory. Science 364, 1082–1086. https://doi.org/10.1126/science.aax0758.

Foster, D.J., and Wilson, M.A. (2007). Hippocampal theta sequences. Hippocampus 17, 1093–1099. https://doi.org/10.1002/hipo.20345.

Gauy, M.M., Lengler, J., Einarsson, H., Meier, F., Weissenberger, F., Yanik, M.F., and Steger, A. (2018). A Hippocampal Model for Behavioral Time Acquisition and Fast Bidirectional Replay of Spatio-Temporal Memory Sequences. Front Neurosci-Switz 12, 961. https://doi.org/10.3389/fnins.2018.00961.

Geiller, T., Vancura, B., Terada, S., Troullinou, E., Chavlis, S., Tsagkatakis, G., Tsakalides, P., Ócsai, K., Poirazi, P., Rózsa, B.J., et al. (2020). Large-Scale 3D Two-Photon Imaging of Molecularly Identified CA1 Interneuron Dynamics in Behaving Mice. Neuron 108, 968–983.e9. https://doi.org/10.1016/j.neuron.2020.09.013.

Geisler, C., Brunel, N., and Wang, X.-J. (2005). Contributions of Intrinsic Membrane Dynamics to Fast Network Oscillations With Irregular Neuronal Discharges. J Neurophysiol 94, 4344–4361. https://doi.org/10.1152/jn.00510.2004.

Gillespie, A.K., Maya, D.A.A., Denovellis, E.L., Liu, D.F., Kastner, D.B., Coulter, M.E., Roumis, D.K., Eden, U.T., and Frank, L.M. (2021). Hippocampal replay reflects specific past experiences rather than a plan for subsequent choice. Neuron https://doi.org/10.1016/j.neuron.2021.07.029.

Girardeau, G., Benchenane, K., Wiener, S.I., Buzsáki, G., and Zugaro, M.B. (2009). Selective suppression of hippocampal ripples impairs spatial memory. Nat Neurosci 12, 1222–1223. https://doi.org/10.1038/nn.2384.

Grienberger, C., Milstein, A.D., Bittner, K.C., Romani, S., and Magee, J.C. (2017). Inhibitory suppression of heterogeneously tuned excitation enhances spatial coding in CA1 place cells. Nat Neurosci 20, 417–426. https://doi.org/10.1038/nn.4486.

Griniasty, M., Tsodyks, M.V., and Amit, D.J. (1993). Conversion of Temporal Correlations Between Stimuli to Spatial Correlations Between Attractors. Neural Comput 5, 1–17. https://doi.org/10.1162/neco.1993.5.1.1.

Grosmark, A.D., and Buzsáki, G. (2016). Diversity in neural firing dynamics supports both rigid and learned hippocampal sequences. Science 351, 1440–1443. https://doi.org/10.1126/science.aad1935.

Gupta, A.S., Meer, M.A.A. van der, Touretzky, D.S., and Redish, A.D. (2010). Hippocampal Replay Is Not a Simple Function of Experience. Neuron 65, 695–705. https://doi.org/10.1016/j.neuron.2010.01.034.

Guzman, S.J., Schlogl, A., Frotscher, M., and Jonas, P. (2016). Synaptic mechanisms of pattern completion in the hippocampal CA3 network. Science 353, 1117–1123. https://doi.org/10.1126/science.aaf1836.

Hangya, B., Li, Y., Muller, R.U., and Czurkó, A. (2010). Complementary spatial firing in place cell–interneuron pairs. J Physiology 588, 4165–4175. https://doi.org/10.1113/jphysiol.2010.194274.

Hines, M.L., Davison, A.P., and Muller, E. (2009). NEURON and Python. Front Neuroinform 3, 1. https://doi.org/10.3389/neuro.11.001.2009.

Hopfield, J.J. (1982). Neural networks and physical systems with emergent collective computational abilities. Proc National Acad Sci 79, 2554–2558. https://doi.org/10.1073/pnas.79.8.2554.

Igata, H., Ikegaya, Y., and Sasaki, T. (2021). Prioritized experience replays on a hippocampal predictive map for learning. Proc National Acad Sci 118, e2011266118. https://doi.org/10.1073/pnas.2011266118.

Itskov, V., Curto, C., Pastalkova, E., and Buzsáki, G. (2011). Cell Assembly Sequences Arising from Spike Threshold Adaptation Keep Track of Time in the Hippocampus. J Neurosci 31, 2828–2834. https://doi.org/10.1523/jneurosci.3773-10.2011.

Izhikevich, E.M. (2007). Dynamical Systems in Neuroscience (MIT Press).

Izhikevich, E.M., and Edelman, G.M. (2008). Large-scale model of mammalian thalamocortical systems. Proc National Acad Sci 105, 3593–3598. https://doi.org/10.1073/pnas.0712231105.

Jadhav, S.P., Kemere, C., German, P.W., and Frank, L.M. (2012). Awake Hippocampal Sharp-Wave Ripples Support Spatial Memory. Science 336, 1454–1458. https://doi.org/10.1126/science.1217230.

Jahnke, S., Timme, M., and Memmesheimer, R.M. (2015). A Unified Dynamic Model for Learning, Replay, and Sharp-Wave/Ripples. J Neurosci 35, 16236–16258. https://doi.org/10.1523/jneurosci.3977-14.2015.

Jensen, O., Idiart, M.A., and Lisman, J.E. (1996). Physiologically realistic formation of autoassociative memory in networks with theta/gamma oscillations: role of fast NMDA channels.

Joo, H.R., and Frank, L.M. (2018). The hippocampal sharp wave–ripple in memory retrieval for immediate use and consolidation. Nat Rev Neurosci 19, 744–757. https://doi.org/10.1038/s41583-018-0077-1.

Káli, S., and Dayan, P. (2000). The Involvement of Recurrent Connections in Area CA3 in Establishing the Properties of Place Fields: a Model. J Neurosci 20, 7463–7477. https://doi.org/10.1523/jneurosci.20-19-07463.2000.

Kang, L., and DeWeese, M.R. (2019). Replay as wavefronts and theta sequences as bump oscillations in a grid cell attractor network. Elife 8, e46351. https://doi.org/10.7554/elife.46351.

Kay, K., Chung, J.E., Sosa, M., Schor, J.S., Karlsson, M.P., Larkin, M.C., Liu, D.F., and Frank, L.M. (2020). Constant Sub-second Cycling between Representations of Possible Futures in the Hippocampus. Cell 180, 552–567.e25. https://doi.org/10.1016/j.cell.2020.01.014.

Lee, I., Griffin, A.L., Zilli, E.A., Eichenbaum, H., and Hasselmo, M.E. (2006). Gradual Translocation of Spatial Correlates of Neuronal Firing in the Hippocampus toward Prospective Reward Locations. Neuron 51, 639–650. https://doi.org/10.1016/j.neuron.2006.06.033.

Levy, W.B. (1989). A Computational Approach to Hippocampal Function. Psychol Learn Motiv 23, 243–305. https://doi.org/10.1016/s0079-7421(08)60113-9.

Lisman, J.E., and Jensen, O. (2013). The theta-gamma neural code. Neuron 77, 1002– 1016. https://doi.org/10.1016/j.neuron.2013.03.007.

Lisman, J.E., Talamini, L.M., and Raffone, A. (2005). Recall of memory sequences by interaction of the dentate and CA3: A revised model of the phase precession. Neural Networks 18, 1191–1201. https://doi.org/10.1016/j.neunet.2005.08.008.

Luczak, A., Barthó, P., Marguet, S.L., Buzsáki, G., and Harris, K.D. (2007). Sequential structure of neocortical spontaneous activity in vivo. Proc National Acad Sci 104, 347– 352. https://doi.org/10.1073/pnas.0605643104.

Lytton, W.W., Seidenstein, A.H., Dura-Bernal, S., McDougal, R.A., Schürmann, F., and Hines, M.L. (2016). Simulation Neurotechnologies for Advancing Brain Research: Parallelizing Large Networks in NEURON. Neural Comput 28, 2063–2090. https://doi.org/10.1162/neco_a_00876.

Malerba, P., and Bazhenov, M. (2019). Circuit mechanisms of hippocampal reactivation during sleep. Neurobiol Learn Mem 160, 98–107. https://doi.org/10.1016/j.nlm.2018.04.018.

Marder, E., and Taylor, A.L. (2011). Multiple models to capture the variability in biological neurons and networks. Nat Neurosci 14, 133–138. https://doi.org/10.1038/nn.2735.

Marr, D. (1971). Simple memory: a theory for archicortex. Philosophical Transactions Royal Soc Lond B Biological Sci 262, 23–81. https://doi.org/10.1098/rstb.1971.0078.

Marshall, L., Henze, D.A., Hirase, H., Leinekugel, X., Dragoi, G., and Buzsáki, G. (2002). Hippocampal Pyramidal Cell–Interneuron Spike Transmission Is Frequency Dependent and Responsible for Place Modulation of Interneuron Discharge. J Neurosci 22, RC197–RC197. https://doi.org/10.1523/jneurosci.22-02-j0001.2002.

McNaughton, B.L., and Morris, R.G.M. (1987). Hippocampal synaptic enhancement and information storage within a distributed memory system. Trends Neurosci 10, 408– 415. https://doi.org/10.1016/0166-2236(87)90011-7.

Mehta, M.R., Lee, A.K., and Wilson, M.A. (2002). Role of experience and oscillations in transforming a rate code into a temporal code. Nature 417, 741–746. https://doi.org/10.1038/nature00807.

Milstein, A.D., Li, Y., Bittner, K.C., Grienberger, C., Soltesz, I., Magee, J.C., and Romani, S. (2020). Bidirectional synaptic plasticity rapidly modifies hippocampal representations. Biorxiv 2020.02.04.934182. https://doi.org/10.1101/2020.02.04.934182.

Okamoto, K., and Ikegaya, Y. (2019). Recurrent connections between CA2 pyramidal cells. Hippocampus 29, 305–312. https://doi.org/10.1002/hipo.23064.

O’Keefe, J., and Conway, D.H. (1978). Hippocampal place units in the freely moving rat: why they fire where they fire. Exp Brain Res 31, 573–590. https://doi.org/10.1007/bf00239813.

O’Keefe, J., and Recce, M.L. (1993). Phase relationship between hippocampal place units and the EEG theta rhythm. Hippocampus 3, 317–330. https://doi.org/10.1002/hipo.450030307.

Ólafsdóttir, H.F., Barry, C., Saleem, A.B., Hassabis, D., and Spiers, H.J. (2015). Hippocampal place cells construct reward related sequences through unexplored space. Elife 4, e06063. https://doi.org/10.7554/elife.06063.

Ólafsdóttir, H.F., Bush, D., and Barry, C. (2018). The Role of Hippocampal Replay in Memory and Planning. Curr Biol 28, R37–R50. https://doi.org/10.1016/j.cub.2017.10.073.

Oliva, A., Fernandez-Ruiz, A., Buzsaki, G., and Berenyi, A. (2016). Role of Hippocampal CA2 Region in Triggering Sharp-Wave Ripples. Neuron 91, 1342–1355. https://doi.org/10.1016/j.neuron.2016.08.008.

O’Neill, J., Senior, T.J., Allen, K., Huxter, J.R., and Csicsvari, J. (2008). Reactivation of experience-dependent cell assembly patterns in the hippocampus. Nat Neurosci 11, 209–215. https://doi.org/10.1038/nn2037.

Pelkey, K.A., Chittajallu, R., Craig, M.T., Tricoire, L., Wester, J.C., and McBain, C.J. (2017). Hippocampal GABAergic Inhibitory Interneurons. Physiol Rev 97, 1619–1747. https://doi.org/10.1152/physrev.00007.2017.

Peters, H.C., Hu, H., Pongs, O., Storm, J.F., and Isbrandt, D. (2005). Conditional transgenic suppression of M channels in mouse brain reveals functions in neuronal excitability, resonance and behavior. Nat Neurosci 8, 51–60. https://doi.org/10.1038/nn1375.

Pfeiffer, B.E. (2020). The content of hippocampal “replay.” Hippocampus 30, 6–18. https://doi.org/10.1002/hipo.22824.

Pfeiffer, B.E., and Foster, D.J. (2015). Autoassociative dynamics in the generation of sequences of hippocampal place cells. Science 349, 180–183. https://doi.org/10.1126/science.aaa9633.

Ramirez-Villegas, J.F., Willeke, K.F., Logothetis, N.K., and Besserve, M. (2018). Dissecting the Synapse- and Frequency-Dependent Network Mechanisms of In Vivo Hippocampal Sharp Wave-Ripples. Neuron 100, 1224–1240 e13. https://doi.org/10.1016/j.neuron.2018.09.041.

Reifenstein, E.T., Khalid, I.B., and Kempter, R. (2021). Synaptic learning rules for sequence learning. Elife 10, e67171. https://doi.org/10.7554/elife.67171.

Renno-Costa, C., Lisman, J.E., and Verschure, P.F. (2014). A signature of attractor dynamics in the CA3 region of the hippocampus. Plos Comput Biol 10, e1003641. https://doi.org/10.1371/journal.pcbi.1003641.

Rennó-Costa, C., Teixeira, D.G., and Soltesz, I. (2019). Regulation of gamma-frequency oscillation by feedforward inhibition: A computational modeling study. Hippocampus 29, 957–970. https://doi.org/10.1002/hipo.23093.

Romani, S., and Tsodyks, M. (2015). Short-term plasticity based network model of place cells dynamics. Hippocampus 25, 94–105. https://doi.org/10.1002/hipo.22355.

Roscow, E.L., Chua, R., Costa, R.P., Jones, M.W., and Lepora, N. (2021). Learning offline: memory replay in biological and artificial reinforcement learning. Trends Neurosci 44, 808–821. https://doi.org/10.1016/j.tins.2021.07.007.

Sasaki, T., Piatti, V.C., Hwaun, E., Ahmadi, S., Lisman, J.E., Leutgeb, S., and Leutgeb, J.K. (2018). DENTATE NETWORK ACTIVITY IS NECESSARY FOR SPATIAL WORKING MEMORY BY SUPPORTING CA3 SHARP-WAVE RIPPLE GENERATION AND PROSPECTIVE FIRING OF CA3 NEURONS. Nat Neurosci 21, 258–269. https://doi.org/10.1038/s41593-017-0061-5.

Scharfman, H.E. (1993). Spiny neurons of area CA3c in rat hippocampal slices have similar electrophysiological characteristics and synaptic responses despite morphological variation. Hippocampus 3, 9–28. https://doi.org/10.1002/hipo.450030103.

Singer, A.C., and Frank, L.M. (2009). Rewarded Outcomes Enhance Reactivation of Experience in the Hippocampus. Neuron 64, 910–921. https://doi.org/10.1016/j.neuron.2009.11.016.

Skaggs, W.E., McNaughton, B.L., Wilson, M.A., and Barnes, C.A. (1996). Theta phase precession in hippocampal neuronal populations and the compression of temporal sequences. Hippocampus 6, 149–172. https://doi.org/10.1002/(sici)1098-1063(1996)6:2<149::aid-hipo6>3.0.co;2-k.

Soltesz, I., and Deschenes, M. (1993). Low- and high-frequency membrane potential oscillations during theta activity in CA1 and CA3 pyramidal neurons of the rat hippocampus under ketamine-xylazine anesthesia. J Neurophysiol 70, 97–116. https://doi.org/10.1152/jn.1993.70.1.97.

Sompolinsky, H., and Kanter, I. (1986). Temporal Association in Asymmetric Neural Networks. Phys Rev Lett 57, 2861–2864. https://doi.org/10.1103/physrevlett.57.2861.

Stark, E., Roux, L., Eichler, R., Senzai, Y., Royer, S., and Buzsaki, G. (2014). Pyramidal cell-interneuron interactions underlie hippocampal ripple oscillations. Neuron 83, 467– 480. https://doi.org/10.1016/j.neuron.2014.06.023.

Stark, E., Roux, L., Eichler, R., and Buzsaki, G. (2015). Local generation of multineuronal spike sequences in the hippocampal CA1 region. Proc National Acad Sci 112, 10521–10526. https://doi.org/10.1073/pnas.1508785112.

Stefanelli, T., Bertollini, C., Luscher, C., Muller, D., and Mendez, P. (2016). Hippocampal Somatostatin Interneurons Control the Size of Neuronal Memory Ensembles. Neuron 89, 1074–1085. https://doi.org/10.1016/j.neuron.2016.01.024.

Stella, F., Baracskay, P., O’Neill, J., and Csicsvari, J. (2019). Hippocampal Reactivation of Random Trajectories Resembling Brownian Diffusion. Neuron 102, 450–461 e7. https://doi.org/10.1016/j.neuron.2019.01.052.

Tremblay, R., Lee, S., and Rudy, B. (2016). GABAergic Interneurons in the Neocortex: From Cellular Properties to Circuits. Neuron 91, 260–292. https://doi.org/10.1016/j.neuron.2016.06.033.

Treves, A. (2004). Computational constraints between retrieving the past and predicting the future, and the CA3-CA1 differentiation. Hippocampus 14, 539–556. https://doi.org/10.1002/hipo.10187.

Treves, A., and Rolls, E.T. (1994). Computational analysis of the role of the hippocampus in memory. Hippocampus 4, 374–391. https://doi.org/10.1002/hipo.450040319.

Tsodyks, M.V., Skaggs, W.E., Sejnowski, T.J., and McNaughton, B.L. (1996). Population dynamics and theta rhythm phase precession of hippocampal place cell firing: A spiking neuron model. Hippocampus 6, 271–280. https://doi.org/10.1002/(sici)1098-1063(1996)6:3<271::aid-hipo5>3.0.co;2-q.

Tukker, J.J., Lasztoczi, B., Katona, L., Roberts, J.D., Pissadaki, E.K., Dalezios, Y., Marton, L., Zhang, L., Klausberger, T., and Somogyi, P. (2013). Distinct dendritic arborization and in vivo firing patterns of parvalbumin-expressing basket cells in the hippocampal area CA3. J Neurosci 33, 6809–6825. https://doi.org/10.1523/jneurosci.5052-12.2013.

Turi, G.F., Li, W.-K., Chavlis, S., Pandi, I., OΓÇÖHare, J., Priestley, J.B., Grosmark, A.D., Liao, Z., Ladow, M., Zhang, J.F., et al. (2019). Vasoactive Intestinal Polypeptide-Expressing Interneurons in the Hippocampus Support Goal-Oriented Spatial Learning. Neuron 101, 1150–1165.e8. https://doi.org/10.1016/j.neuron.2019.01.009.

Wang, X.-J. (2010). Neurophysiological and Computational Principles of Cortical Rhythms in Cognition. Physiol Rev 90, 1195–1268. https://doi.org/10.1152/physrev.00035.2008.

Wilent, W.B., and Nitz, D.A. (2007). Discrete Place Fields of Hippocampal Formation Interneurons. J Neurophysiol 97, 4152–4161. https://doi.org/10.1152/jn.01200.2006.

Wu, X., and Foster, D.J. (2014). Hippocampal Replay Captures the Unique Topological Structure of a Novel Environment. J Neurosci 34, 6459–6469. https://doi.org/10.1523/jneurosci.3414-13.2014.

Ylinen, A., Soltesz, I., Bragin, A., Penttonen, M., Sik, A., and Buzsaki, G. (1995). Intracellular correlates of hippocampal theta rhythm in identified pyramidal cells, granule cells, and basket cells. Hippocampus 5, 78–90. https://doi.org/10.1002/hipo.450050110.

Zaremba, J.D., Diamantopoulou, A., Danielson, N.B., Grosmark, A.D., Kaifosh, P.W., Bowler, J.C., Liao, Z., Sparks, F.T., Gogos, J.A., and Losonczy, A. (2017). Impaired hippocampal place cell dynamics in a mouse model of the 22q11.2 deletion. Nat Neurosci 20, 1612–1623. https://doi.org/10.1038/nn.4634.

Zhang, K., Ginzburg, I., McNaughton, B.L., and Sejnowski, T.J. (1998). Interpreting Neuronal Population Activity by Reconstruction: Unified Framework With Application to Hippocampal Place Cells. J Neurophysiol 79, 1017–1044. https://doi.org/10.1152/jn.1998.79.2.1017.

